# The impact of short-lived controls on the interpretation of lifespan experiments and progress in geroscience

**DOI:** 10.1101/2023.10.08.561459

**Authors:** Kamil Pabis, Diogo Barardo, Jan Gruber, Olga Sirbu, Kumar Selvarajoo, Matt Kaeberlein, Brian K. Kennedy

## Abstract

Although lifespan extension remains the gold standard for assessing interventions hypothesized to impact the biology of aging, there are important limitations to this approach. Our reanalysis of lifespan studies from multiple sources suggests that the use of short-lived control cohorts tends to exaggerate the relative efficacy of putative longevity interventions. Moreover, due to the high cost and long timeframes of mouse studies, it is rare that a particular longevity intervention will be independently replicated by multiple groups.

To facilitate identification of successful interventions, we propose an alternative approach. The level of confidence we can have in an intervention is proportional to the degree of lifespan extension above the strain- and species-specific upper limit of lifespan, which we can estimate from comparison to historical controls. In the absence of independent replication, a putative mouse longevity intervention should only be considered with high confidence when control lifespans are close to 900 days or if the final lifespan of the treated group is considerably above 900 days. Using this “900-day rule” we identified several candidate interventions from the literature that merit follow-up studies.

## Introduction

It is an open secret within the field of geroscience research that short-lived and metabolically unhealthy control animals can complicate the interpretation of lifespan studies. In addition, mouse lifespan studies are often small, limited to one sex and fail to report potential confounding factors. Multiple authors have pointed out these problems and recommended steps to alleviate them (**Spindler 2012, Ladiges et al. 2009, Bronwen et al. 2010, Bischoff and Volynets 2016**).

Incorporating many of these suggestions for optimal mouse husbandry and avoiding pitfalls of other lifespan studies, the rigorous National Institute of Aging Interventions Testing Program (ITP) has become a gold-standard for mouse longevity studies (**Nadon et al. 2017**). In the ITP, studies are performed on both sexes, with large sample sizes and across three different centers to address idiosyncratic issues of mouse husbandry. Furthermore, the UM-HET3 mice used by the ITP are relatively long-lived compared to most inbred mouse strains and genetically heterogenous, thereby reducing the likelihood that mice die of strain-specific pathologies, a factor that may confound lifespan data.

A majority of compounds tested by the ITP have not been previously published to extend lifespan in mice, thus we lack a “ground truth” for their expected effect size. Notably, however, the ITP has failed to replicate published lifespan extension for several compounds such as metformin (**Strong et al. 2016**), resveratrol (**Strong et al. 2013**) and nicotinamide riboside (**Harrison et al. 2021**), raising significant concerns about the overall quality of published mouse longevity data.

Although differences in genetic background, age of treatment onset, husbandry, and dosing between the original study and the ITP cohorts may explain the failure to replicate, another potential factor is methodological rigor. For example, many of the ITP-tested compounds that were supported by positive published data had already produced inconsistent results in earlier studies, e.g. aspirin (**Hochschild 1973**), or only minimal lifespan extension (<5%), e.g. nicotinamide variants (**Zhang et al. 2016**) and metformin (**Martin-Montalvo et al. 2013**). In other cases, compounds were predominantly tested in short-lived and/or unhealthy controls, e.g. resveratrol (**Baur et al. 2006**) and curcumin (**Kitani et al. 2004**). Avoiding the above-mentioned experimental shortcomings already at the study conception stage could reduce the amount of time and money spent on failed replication efforts and follow-up studies, thereby improving reproducibility of mouse research and accelerating progress towards truly geroprotective compounds.

In this manuscript, we reanalyze data from CR studies performed in multiple species, from the ITP and from large mouse lifespan studies with a particular focus on control lifespan as one potential explanation for inflated effect sizes and lack of reproducibility. We show that both statistical and biological causes exaggerate the benefits of interventions tested against short-lived controls. As a solution, we propose the use of long-lived controls in mouse studies which should reach a lifespan of around 900 ±50 days, or the comparison to appropriate historical controls, and we term this the “900-day rule”. Finally, applying this new rule, we compare reported interventions to uncover the most promising candidates for follow-up studies.

## RESULTS

### Why do short-lived controls matter? The metformin case-study

We will discuss metformin as an illustrative example where, even prior to ITP testing, discrepancies were apparent in the literature. Early work in very short-lived mice (lifespan<300 days) suggested that biguanides like metformin and phenformin could extend lifespan and prevent cancer (**Anisimov et al. 2003, Anisimov et al. 2005**). It was not until 10 years later that metformin was tested in healthier mice. Since then, many studies have tested the effects of metformin with results ranging from small lifespan extension (**Martin-Montalvo et al. 2013**), over no effect (**Strong et al. 2016, Alfaras et al. 2017**) to a small reduction (**Zhu et al. 2021**).

Altogether, a recent meta-analysis suggested that metformin does not significantly extend lifespan in mice (**Parish and Swindell 2022**). Metformin seemed to work less well in studies involving longer-lived mice, like in the ITP. Using data from the recent meta-analysis we explored this possibility in more detail. When we plot the absolute (**Fig. S1A**) or relative (**Fig. S1B**) change in median lifespan in metformin studies against the lifespan of control mice we notice a striking negative correlation. This correlation was not sensitive to the inclusion or exclusion of specific datasets. We saw the same kind of relationship when we analyzed results from the ITP separately (**Fig. S1C, D**), when we excluded the ITP data from the meta-analysis (**Fig. S1E, F**) or when we excluded the studies by Anisimov et al. from the analysis due to their short lifespan (**Fig. S1G, H**). Importantly, since high doses of metformin are toxic, we confirmed that similar results are also seen in studies with lower doses of the drug (<1000ppm, **Fig. S2A, B**). These findings led us to revisit the importance of control lifespans as a determinant for the reproducibility and robustness of mouse lifespan studies.

### Short-lived strains within a species respond more favorably to lifespan-extending interventions

It is possible that the inverse relationship we saw between control lifespan and the effects of metformin was confounded by the differences in mouse strain, drug dose or husbandry conditions between studies. Therefore, to mitigate this problem we searched the literature for studies that maintained consistent husbandry conditions and subjected cohorts with varying genetic backgrounds to a fixed drug or longevity treatment.

Such study designs are rare and none have been undertaken with lifespan extending drugs. Therefore, instead we re-analyzed the raw data from four large studies that imposed caloric restriction (CR) in yeast (**Schleit et al. 2013**), worms (**Snoek et al. 2019**), flies (**Jin et al. 2020**) and in recombinant inbred ILSXISS mice (**Liao et al. 2010, Rikke et al. 2010, Unnikrishnan et al. 2021**) with differing lifespan. In all these studies differences in control lifespan are primarily due to genetic determinants because the cohorts were kept under identical conditions in the same lab and subjected to the same degree of CR.

We find that cohorts with higher lifespan of control animals (“control LS” in short) show less lifespan extension with CR and other interventions (**Fig. 1-3, Table S1-3**) and that many longevity promoting interventions merely move the median lifespan closer to the strain-specific optimum and do not extend it further (idealized example shown in **Fig. 1A, B**). This becomes more obvious in the case of CR when we plot the fold-change in lifespan for the top 10% of longest-lived strains and the bottom 10% of the shortest-lived strains in each of the four species considered. Indeed, across all the species CR was unable to extend the lifespan of the 10% longest-lived strains (**Fig. 1C**). Instead, control lifespan appears to mediate the effect of CR on lifespan extension, explaining more variability in lifespan extension in long-lived species like mice as compared to short-lived yeast (**Fig. 1D**).

**Figure 1.**
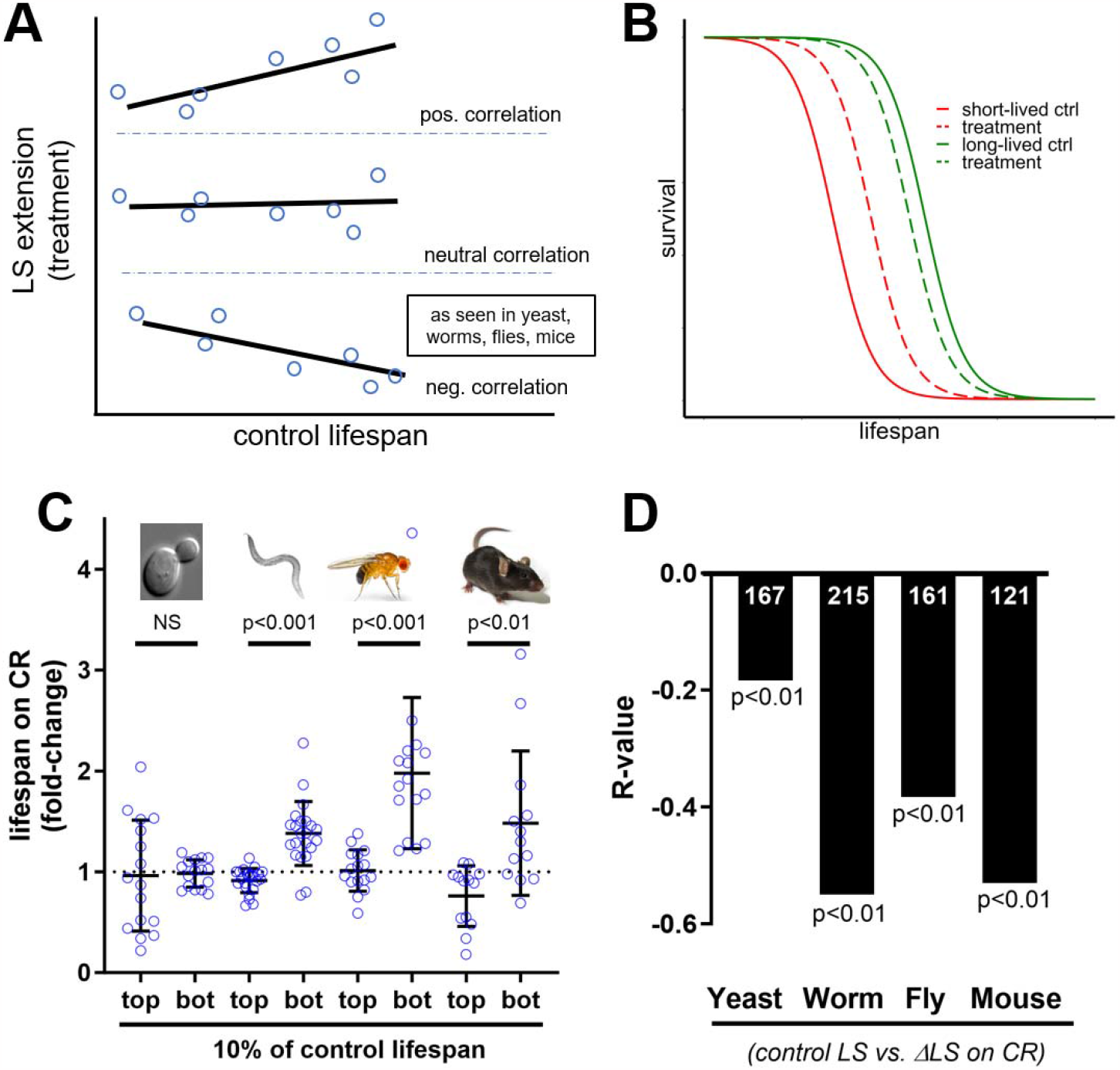
Longer-lived strains within species respond less favorably to caloric restriction (CR) (A) The three possible correlation patterns between control lifespan (LS) and the effect of treatments on LS extension: positive relation (top panel), neutral relation (mid) and negative relation (bottom, consistent with observed data). (B) The inverse correlation in (A) can be explained when a treatment leads to LS extension relative to short-lived controls (the red line moves towards the dashed red line) and LS shortening or no effect compared to long-lived controls (the green lines moves towards the dashed green line). (C) Fold-change in LS extension under caloric restriction (CR) for the top 10% longest-lived strains (“top”) and the bottom 10% shortest-lived strains (“bot”) in each species. The longest-lived worm, fly and mouse strains show no LS extension under CR, whereas the shortest-lived strains do. This pattern is not evident in yeast. P-values based on T-test for unequal variances. (D) The correlation, expressed as absolute R-value, between control LS and LS extension under CR for different species shows a negative trend, where more negative values mean that long-lived strains within this species respond less favorably to CR. Sample sizes are indicated in a white font (number of cohorts). Data for yeast is from **Schleit et al. (2013)**, for worms from worms (**Snoek et al. 2019**), for flies from **Jin et al. (2020)** and **Wilson et al (2020)**, and for mice from (**Liao et al. 2010, Rikke et al. 2010, Unnikrishnan et al. 2021**).

### Short control lifespans exaggerate the benefits of CR due to a mix of technical and biological causes

Next, we reanalyzed the mouse data from **Fig. 1** in more detail. When we plot control lifespan against lifespan extension by CR we see a negative relationship in female (**Fig. 2A**) and male mice (**Fig. 2B**) individually, and in the pooled dataset (**Fig. 1D**).

**Figure 2.**
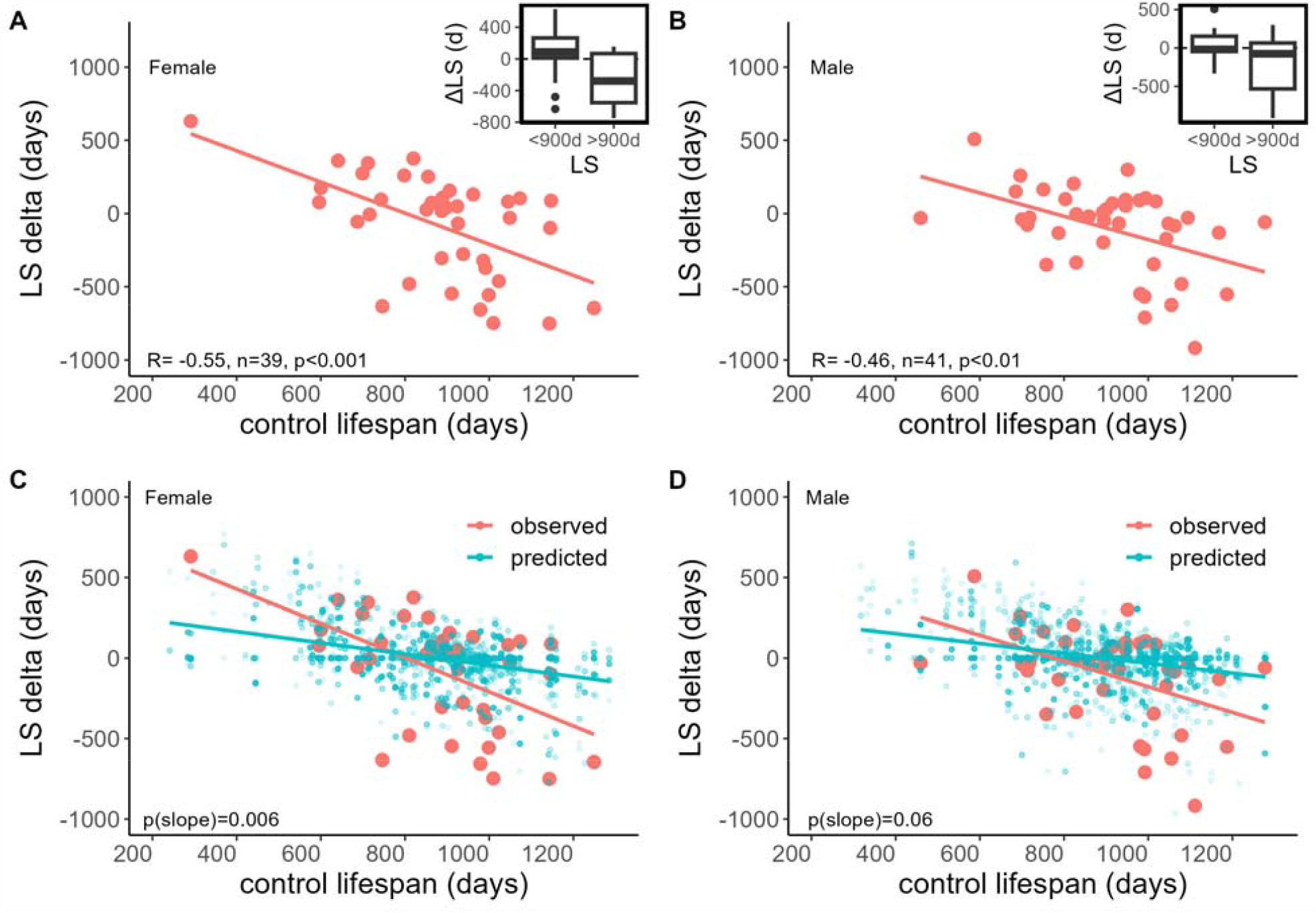
Long-lived female and male ILSXISS strains respond less favorably to caloric restriction. Lifespan (LS) of female (A) and male (B) control mice from different strains on the X-axis (pink dots) plotted against the absolute change in lifespan with caloric restriction (CR) on the Y-axis when imposed in the respective strain (Δlifespan CR). Mouse cohorts with a lifespan of <900 days benefit from CR whereas mice with a lifespan of >900 days do not (see the insert). To test whether regression to the mean can explain exaggerated benefits in short-lived mice we resampled quasi-lifespan experiments from the control population. The resampled synthetic data (blue) is shown for female (C) and male (D) mice with the observed datapoints overlaid (pink). Figures based on data deposited in the Mouse Phenome Database which is comprised of a subset from **Rikke et al. (2010)** and **Liao et al. (2010)**.

Before continuing our analysis of this dataset, we sought to address concerns that the small group sizes in these studies preclude reliable determination of lifespan (**Mattson 2010**). If this were the case, then measurements between different labs should produce mutually inconsistent lifespan data. However, using data from three different experiments (**Liao et al. 2010, Rikke et al. 2010, Unnikrishnan et al. 2021**), we were able to confirm that the strain-specific lifespans were significantly correlated between these studies (**Fig. S3A, C**). The CR response was not significantly correlated between studies (**Fig. S3B, D**).

Since the same strains have a similar lifespan across studies, this indicates that genetic differences underlie lifespan differences between strains in these studies. Importantly, this makes it less likely that the effects of CR are fully explained by regression to the mean, which may arise due to stochastic sampling and give rise to an apparent negative relationship between control lifespan and intervention lifespan (**Garratt et al. 2017**). We tested this by resampling from the control population to generate both control and treated groups. Any negative relationship between control lifespan and lifespan extension based on these resampled values should be purely spurious.

However, consistent with a biological effect on top of regression to the mean, the observed regression line had a significantly more negative slope than the resampled line in female mice (**Fig. 2C**), with a trend in males (p=0.06, **Fig. 2D**). Therefore, long-lived ILSXISS strains responded less favorably to CR than expected based on regression to the mean effects. Although below we will focus on mice only, it is reassuring that the invertebrate data is in agreement. In studies of calorie restricted flies, lifespan was extended (**Fig. S4A, B**), while long-lived fly strains responded less favorably to CR than expected based on regression to the mean effects (**Fig. S4C**). Similarly, restricted worms also lived longer (**Fig. S4D, E**) and long-lived strains responded less favorably to CR than expected (**Fig. S4F**).

Further underscoring the consistency of our findings, we saw a negative correlation in all three ILSXISS studies published several years apart (**Fig. S5**), and after outlier removal (R^female^= -0.47 and R^male^= -0.43; p<0.01). Although the authors attributed part of the CR response in these mice to fat maintenance and changes in body temperature (**Liao et al. 2011**), this does not explain our results. Ad libitum lifespan remained a significant predictor of CR response when controlling for fat loss (p<0.01, n=71) and was borderline significant when controlling for change in body temperature in a smaller subset of mice (p=0.051, n=26). To the contrary, our results suggest that ad libitum lifespan can, to some extent, mediate the link between fat maintenance and CR response (**Fig. S6**).

### Short control lifespans exaggerate the benefits of interventions reported in meta-analyses

To assess whether the above findings can be replicated outside of the context of CR, and when pooling highly heterogenous data, we reanalyzed several, large meta-analytic datasets (**Swindell et al. 2012, Barardo et al. 2017, Garratt et al. 2017, de Magalhães et al. 2018**).

First, we reanalyzed lifespan data from a meta-analysis of CR studies by **Swindell et al. (2012)**, after excluding the ILSXISS data, to test whether studies with longer-lived controls showed smaller lifespan extension after CR. No correlation was seen in mice between control lifespan and lifespan extension (**Fig. S7A**), while there was a small negative correlation in rats (**Fig. S7B**). Interestingly, we did see a significant correlation in mice when we looked at the single largest dataset in this meta-analysis (n=15; **Fig. S8A, B**), suggesting that differences in husbandry conditions between studies could mask an effect of control lifespan.

Next, we reanalyzed mouse longevity interventions from the DrugAge database (**Barardo et al. 2017**). Although our data extraction strategy was different from the original publication, since we focused on absolute rather than relative lifespans, our results are nonetheless in good agreement with the reported lifespan extension in DrugAge (**Fig. S9**). No significant negative correlation was observed between control lifespans and drug-induced lifespan extension (R= -0.09, n=147).

As was the case for CR studies, the single largest dataset in DrugAge (n=22) revealed a strong negative correlation between control lifespan and treatment effect. **Schroeder and Mitchener (1975)** tested the impact of different metals on the longevity of male and female Swiss mice across multiple experiments with varying control lifespans. In this dataset we found a significant, steep and negative correlation between control lifespan and treatment effect (**Fig. S10A, B**).

Two meta-analyses of genetic interventions also both found evidence for an impact of control lifespan on the lifespan extension in various mutant mouse models. In our re-analysis of **Garratt et al. (2017)** we found that both longer-lived IGF1/IRS mutants (**Fig. S11A, B**) and GH dwarfs (**Fig. S11C, D**) were less likely to show lifespan extension. Similarly, the meta-analysis by **de Magalhães et al. (2018)** found that control lifespans significantly influenced the lifespan extending effects of genetic interventions (R= -0.55, n=33).

All in all, the strong negative relationship between control lifespan and treatment effect seen in large, highly controlled studies with multiple cohorts (**Fig. 2; Fig. S8A, B; Fig. S10A, B**) contrasts with a weaker relationship in meta-analyses. This suggests that between-study variability could mask the effects of control lifespan on experimental lifespan extension (**Table S4**).

### Short control lifespans exaggerate the benefits of drugs tested in the Interventions Testing Program via “regression to the mean”

Since large heterogeneity in husbandry and interventions between experiments could mask the effect of control lifespan in meta-analysis, we searched for studies that tested different interventions under more comparable conditions. The only large study with consistent husbandry conditions that we identified was the ITP (**Nadon et al. 2017**).

The ITP dataset we analyze includes raw data for 68 drugs tested across 3 study sites. Since drugs are usually tested in both sexes, this yields 395 conditions in total, where a condition is defined as a particular combination of drug x gender x testing site.

Using the aggregated summary data, we again found a negative correlation between control lifespan and treatment effect in the ITP (R= -0.22, p<0.05, n=132; **Fig. S12**). This correlation becomes even more apparent when treating the results from each testing site as independent experiments (R= -0.27, p<0.0001, n=395; **Fig. 3**). The latter analysis may be more appropriate than one considering the aggregate data, as there are large differences in the lifespan of mice between testing sites.

**Fig. 3.**
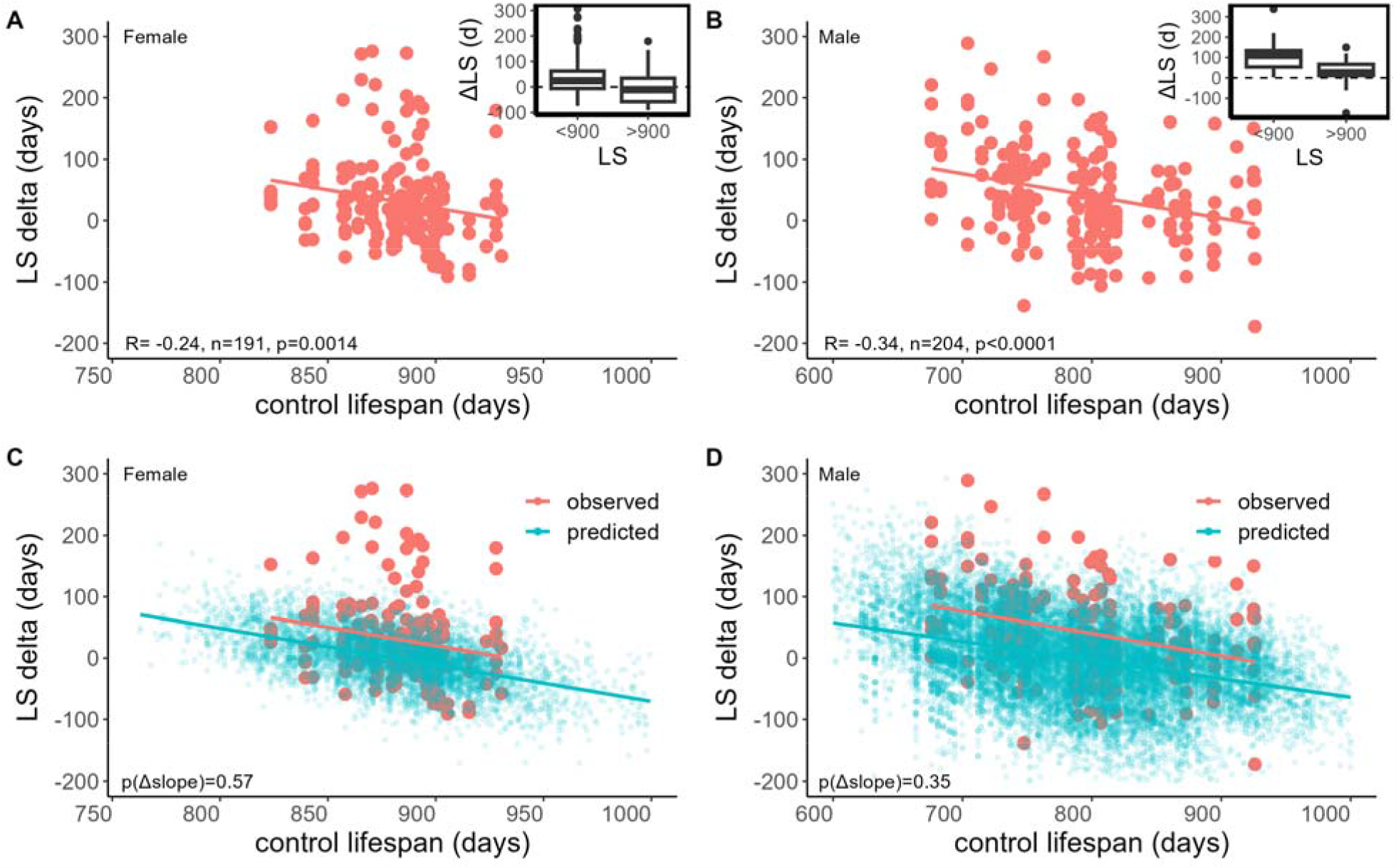
Longer-lived cohorts of UM-HET3 mice show less pronounced lifespan extension in the interventions testing program (ITP) Lifespan (LS) of female (A) and male (B) control mice in the ITP study (pink dots) plotted against the change in lifespan with drug treatments on the Y-axis (Δlifespan in days). Each point corresponds to a unique combination of drug x gender x testing site. Mouse cohorts with a lifespan of <900 days benefit more from drug treatments than do mice with a lifespan of >900 days (see the insert). To test whether regression to the mean can explain exaggerated benefits in short-lived mice we resampled quasi-lifespan experiments from the control population. The resampled synthetic data (blue) is shown for female (C) and male (D) mice with the observed datapoints overlaid (pink). P-value in (A) and (B) based on a linear mixed effects model considering cohort and control lifespan.

Cohorts of longer-lived UM-HET3 mice showed less lifespan extension in response to various treatments whether lifespan extension was defined in absolute (**Fig. S13A**) or relative terms (**Fig. S13B**). Importantly, a significant negative correlation between control lifespan and treatment effect was seen in both females (**Fig. 3A**) and males (**Fig. 3B**), and across multiple testing sites, specifically the University of Texas Health Science Center for both sexes and the Jackson Laboratory for males (**Table S5**). However, our resampling analysis indicated that this effect was largely due to regression to the mean since the observed and the resampled regression line were almost parallel (**Fig. 3C, D**).

Arguably, the results in **Fig. 3** may be an imprecise estimate of the true relationship because each treatment contributes only a few datapoints to the correlation. However, the ITP also provides a unique opportunity to address this issue. Since each drug was tested across three study sites with different control lifespan, we can perform a Spearman correlation analysis for every drug. We find a negative correlation between control lifespan and treatment effect in the pooled analysis for 51 out of 68 drugs tested (75%, p<0.0001; p-value by permutation). Split by gender, we find a negative correlation between control lifespan and treatment effect for 52 out of 68 drugs in males (76%, p<0.0001) and 40 out of 68 drugs in females (59%, p=0.053).

### Control lifespans explain differential sex effects in the Interventions Testing Program

In the previous section (**Figures 3A, B**), we observed a stronger negative correlation in male mice, which may partially explain some of the sexually dimorphic drug responses in the ITP. Indeed, control males are shorter-lived than females (median 798 vs 882 days, p<0.0001) and also respond better to interventions (+38 vs 27 days, p=0.10). The pooled data, however, underestimates these sex differences. Since rapamycin shows elevated blood levels in female mice (**Miller et al. 2014**) and was tested more frequently than any other drug in the ITP (making up 16% of all interventions), this will increase the apparent lifespan extension seen in female mice (**Fig. S14A**). Thus, if we only consider one unique result per drug, male mice respond much better than females (+32 vs 7 days lifespan extension, p<0.01; **Table 1**) with 66% of the drugs producing higher lifespan extension in ITP males (**Fig. S14B**) and no similar sex dimorphic benefits observed in DrugAge (**Fig. S14C**).

**Table 1.** Lifespan extension in male and female mice (different subsets of the ITP) We calculated the mean lifespan (LS) extension for male and female mice in the ITP after excluding redundant results from drugs that were tested multiple times. Males benefit more from longevity extending interventions in the ITP whether we analyze the pooled data (“pooled”) or treat each study site as an independent experiment (“cohort-level”). The p-value is for the difference between the lifespan extension of male and female cohorts (paired T-test).

|  |  | male | female |  |
| --- | --- | --- | --- | --- |
| Subset | N | LS-extension (days) |  | p-value |
| pooled | 41 | 32.1 | 6.7 | 0.001 |
| cohort-level | 123 | 25.3 | 9.3 | 0.030 |

Interestingly, the significant male advantage we observed was partly driven by a better response of male cohorts at the University of Texas (**Fig. S14D**), where male mice are particularly short-lived compared to females (**Table 2**). Significant findings in males at this site were almost two times more common than at the Jackson Laboratory or the University of Michigan sites and more common than in female mice at the same site (**Table S6**).

**Table 2.** Control lifespans at the three testing sites of the ITP. Lifespans (LS) in days from all control cohorts across the three study sites of the ITP (TJL, UM and UT). For male cohorts, the average increase of LS after drug treatment (ΔLS) is highest at the UT site where the average LS of controls is shortest, whereas this is not the case for female cohorts. LS are a mean of median LS reported for each study year. TJL=The Jackson Laboratory, UM=University of Michigan, UT=University of Texas Health Science Center.

|  | The Jackson Laboratory |  | University of Michigan |  | University of Texas |  |
| --- | --- | --- | --- | --- | --- | --- |
|  | LS (days) | ΔLS (drug) | LS (days) | ΔLS (drug) | LS (days) | ΔLS (drug) |
| male | 782 | 31.3 | 857 | 37.2 | 753 | 66.3 |
| female | 887 | 45.2 | 887 | 19.7 | 872 | 32.6 |

If we look at individual treatments that significantly extended lifespan in either gender, we can see a pattern consistent with a male advantage. As discussed elsewhere, acarbose benefits males more, while the opposite is seen for rapamycin (**Harrison et al. 2014, Miller et al. 2013**). However, most other drugs for which lifespan extension has been reported clearly benefit males, e.g. NDGA (C2004: +90d difference male vs female), 17α-Estradiol (C2009: +64d), canagliflozin (C2016: +98d), aspirin (C2004: +100d), protandim (C2011: +58d). Similarly, while captopril and glycine benefited both sexes the benefit was larger in males for captoptril (C2017: +55d) and glycine (C2014: +19d). In contrast, the only drug that reached significance in females but not males was 1,3-butanediol, although the absolute lifespan extension was still larger in males (C2017: +54d).

### The “900-day rule” defines a lifespan gold standard for mouse lifespan studies

Since C57BL/6 and UM-HET3 are currently the most important mouse strains in aging research, we provide normative median lifespans for these and compare them with other strains.

As demonstrated by **Austad (2011)**, using only studies providing lifespan for both male and female mice, there is no clear sex difference in lifespan of C57BL/6 mice (**Fig. S15**). However, we found that median lifespans of C57BL/6 mice are quite variable and depend on the dataset used. For males, lifespan range from 779 days to 861 days and for females from 720 days to 910 days in different meta-analyses (**Fig. 4**). In addition, we investigated whether there are lifespan differences between the Jax and Nia substrains of C57BL/6 mice, which have been shown to differ in some important traits (**Mekada and Yoshiki 2021**). Although, most of the studies used the C57BL/6J substrain the few studies using C57BL/6Nia reported comparable lifespans (**Fig. S16**).

**Figure 4.**
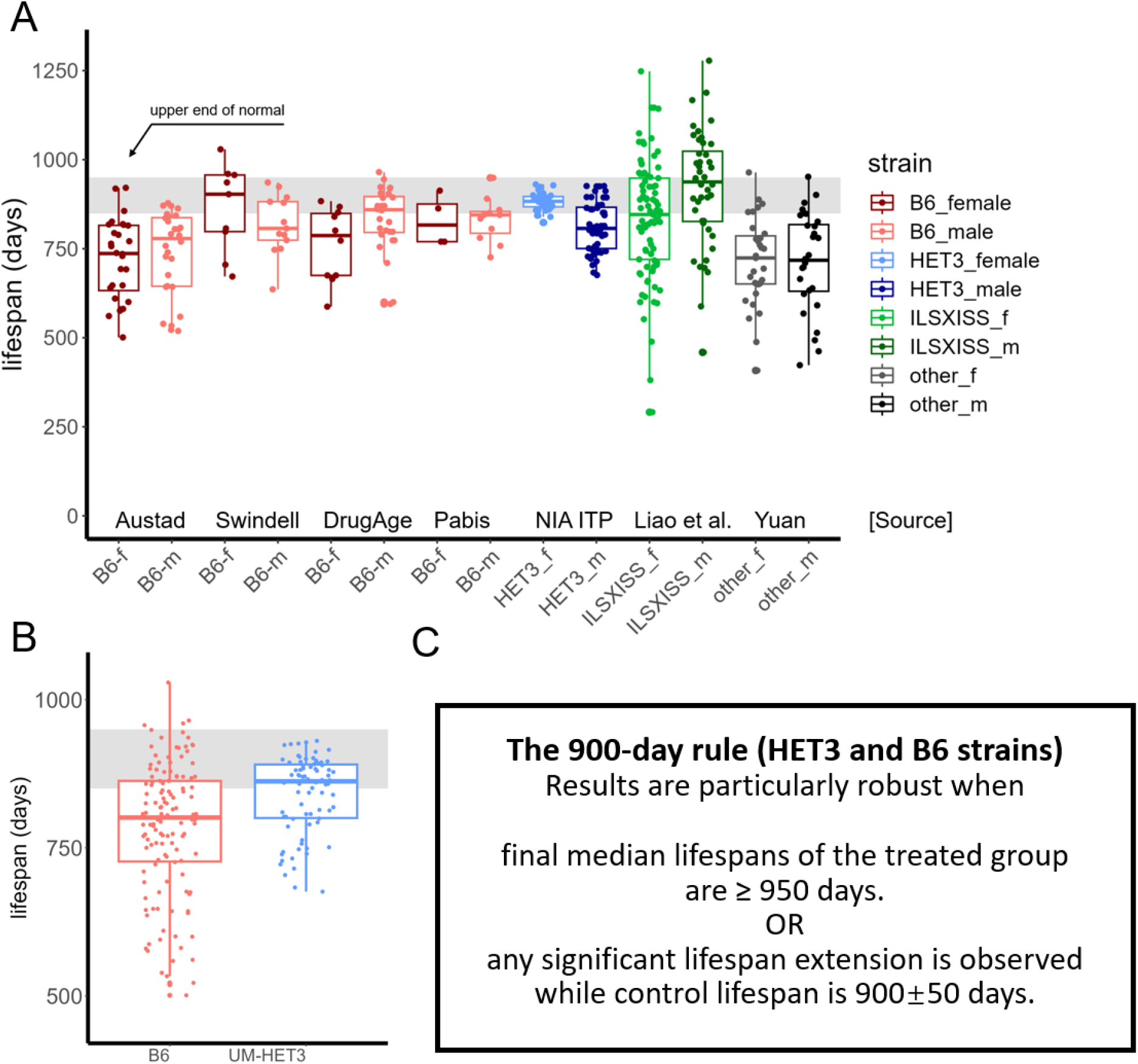
Healthy inbred and hybrid mouse strains live close to 900 days. A) Under normal conditions, healthy control mice live close to 900 days. From left to right, lifespans for female (f) and male (m) C57BL/6 (B6) mouse cohorts from **Austad (2011), Swindell et al. (2012)**, DrugAge (**Barardo et al. 2017**) and from our own analysis. This is followed by lifespans for UM-HET3 (HET3) mouse cohorts tested by the ITP. Finally, for comparison we show data from the ILSXISS inbred panel (**Liao et al. 2010, Rikke et al. 2010, Unnikrishnan et al. 2021)** and from **Yuan et al. (2012)**. Lifespans in the original datasets are either mean or median, depending on data-availability. The interval between 850 and 950 days is indicated with a shaded area. Boxplots show median ± 95% CI. B) Pooling all the B6 and HET3 data from (A) it becomes more obvious that 900 days represents the upper end of normal for these strains and few published cohorts using wildtype mice showed median lifespans considerably above that value. The here reported values can serve as historical controls for comparison purposes. C) Based on these findings, the 900-day rule can be phrased in two ways. 1. It would be unusual to observe median lifespans considerably above 900 days in a mouse experiment, hence lifespan extension above 950 days - to allow for a buffer - compared to historical controls indicates that the given treatment shows robust lifespan extension, 2. If the controls are long-lived, i.e. 900±50 days, then any significant lifespan extension observed is more likely to be robust and not due to amelioration of premature death.

In the case of UM-HET3 mice from the ITP cohorts, males appear to be shorter-lived. Female median lifespan was 883 days and male median lifespan was 800 days. As discussed before, however, male lifespans are dependent on the testing site (**Table 2**). Males at the University of Michigan show a median lifespan of 857 days not too different from female UM-HET3 mice.

For comparison, we plotted lifespans from two other large datasets. Mice from the ILSXISS inbred panel are very long-lived, albeit with a lot of variability. These strains had a median lifespan of around 882 days, with a lifespan of 938 days for males and 835 days for females (**Table S7**). In contrast, the 32 commonly used inbred strains whose lifespan was reported by **Yuan et al. (2012)** are rather short-lived with a median lifespan of 721 days, and, with a lifespan of 718 days for males and 724 days for females. However, in this study, C57BL/6J was among the longest-lived strains with males reaching a median lifespan of 901 days and females of 866 days.

There are several reasons to suggest that researchers should work with the longest-lived mice they can. Not only have we documented exaggerated lifespan extending effects in experiments with shorter-lived controls (**Fig. 3**). Moreover, it could be argued that long-lived strains are a more faithful model for human physiology and longevity given the exceptionally long lifespans of humans compared to other animals (**Buffenstein 2009**).

Thus, we propose the “900-day rule” for mouse lifespan experiments, which is easy to remember and sufficiently accurate to be useful to editors, reviewers, scientists and lay readers alike. Most healthy inbred or hybrid strains should have a median lifespan of close to 900 days (± 50 days). Since the normative lifespans we presented here are likely a lower bound for the true strain-specific lifespan of these animals, due to husbandry issues, we believe that C57BL/6, UM-HET3 and some other strains are well capable of such lifespans (**Table S7**). Based on the 900-day rule we define treatments that extend the lifespan of short-lived cohorts as “longevity-normalizing”, whereas those that work in long-lived cohorts are “longevity-extending”.

Importantly, without an appropriately long-lived control, it is impossible to attribute lifespan extension to effects on biological aging since the tested intervention could be simply offsetting idiosyncratic health issues. However, in the absence of a long-lived within-study control these values (**Fig. 4; Table S8**) can serve as a historical control. Interventions that result in median cohort lifespans well above 900 days in mice should be taken seriously independent of the within study controls (**Fig. 4C**). Conversely, even large lifespan increases against a short-lived background may be artefactual. As a corollary, the use of percentage increase in lifespan should be discouraged because it fails to capture, and indeed can often conceal, essential information about control lifespan.

While plausible, the question remains if such a simple rule can successfully predict robust interventions? To test this, we asked whether interventions identified in DrugAge that passed the 900-day rule would be more likely to extend lifespan in the ITP than interventions which failed the rule. Although the available data for compounds found in both datasets is limited, NDGA and rapamycin were the only intervention that showed lifespan extension in long-lived DrugAge cohorts and it were also relatively successful interventions in the ITP (**Table S9**).

### Re-ranking of interventions using meta-analysis and absolute lifespans

Using the 900-day rule we identified 19 interventional groups in the ITP that meet our criteria in at least one cohort (**Table S10**). As expected, these included acarbose, rapamycin and 17-α-estradiol but also other compounds like glycine or captopril. In total 10 unique compounds met the cut-offs. However, when data from all three cohorts was pooled, no interventions met our criteria except rapamycin, and rapamycin combinations, in female mice (**Table S11**). This suggests that few compounds consistently increase lifespan across multiple cohorts of long-lived UM-HET3 mice.

Nevertheless, compounds that are beneficial in a few cohorts may still be worth exploring. To account for cohort lifespan variation in a more fine-grained way and identify such compounds, we constructed a linear regression model that considers the sex, treatment, site and control lifespan of a cohort. We then searched for compounds that produce 50 days more lifespan extension than predicted. This identified 70 interventional groups comprising 29 unique compounds. Although this analysis broadly agrees with the findings of the ITP, which are based on log-rank test statistics, we identify several additional compounds that could be promising for lifespan extension (**Table S12**). For example, our analysis suggests that inhibition of angiotensin converting enzyme is beneficial to mouse lifespan since both captopril and enalapril led to higher-than-expected lifespan extension in some cohorts, although the benefits were most pronounced in males for captopril (**Fig. S17A, B**) and exclusively seen in males for enalapril (**Fig. S17C, D**).

Using a similar approach as in **Table S12** we re-evaluated the efficacy of rapamycin combination treatments in the ITP. These combinations were tested without a rapamycin control during the same year and thus necessitate a comparison with historical controls. When we rank compounds by lifespan extension^actual-predicted^ in each cohort we find that rapamycin (14 ppm) combined with either acarbose or metformin leads to higher lifespan extension than do most other rapamycin groups (**Table S13**). The combined rapamycin groups also outperform rapamycin-only groups when we rank all interventions by the median lifespan of the treated group (**Fig. S18**). When we limited the comparison to the closest matched rapamycin groups (14 ppm started at 9-months), the combination of metformin and rapamycin led to significantly higher lifespan extension than just rapamycin alone (**Fig. S19A**) and combination treated animals were longer-lived in absolute terms than rapamycin treated animals (**Fig. S19B**). When we plot the full survival curves compared to historical controls, the lifespan extension is most pronounced in male mice (**Fig. S20A, C**) rather than female mice (**Fig. S20B, D**).

Next, in our reanalysis of DrugAge we found 14 datasets comprising 12 different compounds that met the 900-day rule (**Fig. 5A, Table S14**). Interestingly, this set included three drugs that reduce heart rate, i.e. the two beta-blockers, metoprolol and nebivolol, and ivabradine.

**Figure 5.**
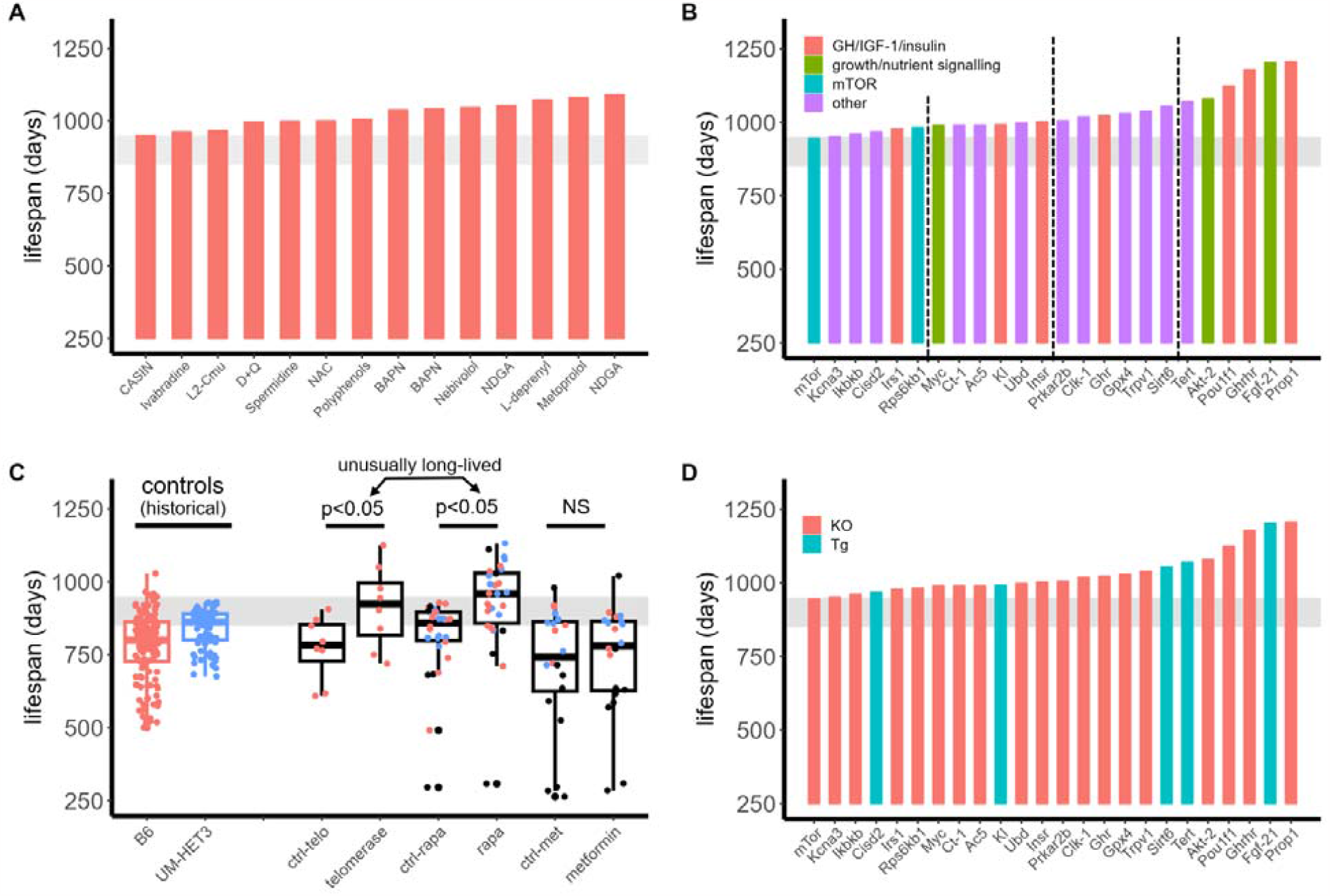
Certain drugs and genetic interventions extend mouse lifespan compared to historical controls. For this figure any intervention producing a final median lifespan of ≥ 950 days was considered to pass the 900-day rule. A) 14 different cohorts with 12 unique compounds from DrugAge (Barardo et al. 2017) pass the 900-day rule. Abbreviations: D+Q = dasatinib + quercetin, NDGA = nordihydroguaiaretic acid, NAC = N-acetyl-L-cysteine, BAPN = beta-Aminopropionitrile fumarate, CASIN = Cdc42 inhibitor, L2-Cmu = IGF-1R mAb. B) 23 different genetic interventions reported in GenAge (**Tacutu et al. 2018**) pass the 900-day rule. Although the mTOR hypomorphic strain failed the 900-day rule by a small margin (treated LS of 945 days) it was included as the 24th intervention due to prior plausibility. C) Mice treated with rapamycin (rapa) or subjected to telomerase activation live longer than most historical controls. From left to right, lifespans for C57BL/6 (B6) and UM-HET3 mouse cohorts of both sexes (n=131 and 78, respectively, based on the data in Fig. 4) used as historical controls. Followed by data from telomerase induced cohorts (n=8 per group), rapamycin treated cohorts (n=30, data from **Selvarani et al. 2021**), and metformin treated cohorts (n=20, **Parish and Swindell 2022**) with the respective control (ctrl) and treated arm. The telomerase data includes studies using viral vectors and transgenic mice. The interval between 850 and 950 days is indicated with a shaded area. Boxplots show median ± 95% CI. P-values based on paired T-test. D) The majority of interventions that robustly extend lifespan in GenAge are gene knock-outs (KO), whereas only few transgenic (Tg) mouse models were reported to extend lifespan.

Having shown that the 900-day rule can inform the interpretation of mouse lifespan studies using pharmacologic interventions, we extended our analysis to genetic studies reported in GenAge (**Tacutu et al. 2018**). We identified 24 out of 136 longevity genes that extended lifespan in studies with long-lived control mice (**Table S15**). These fell into four major categories: mTOR signalling, growth signalling, GH/IGF-1/Insulin-axis and diverse other pathways (e.g. telomerase, DNA repair or inflammation).

To narrow down the top genes we ranked the 24 candidates by the absolute lifespan of the intervention group and excluded interventions that led to lifespans of <950 days (**Fig. 5A**). The longest-lived animals were knock-outs in the growth hormone pathway (Ghrhr, Prop1, Pou1f1). Several other genes were also associated with exceptionally long lifespans in at least one studied cohort. This includes the overexpression of genes involved in DNA repair (Sirt6), telomere extension (Tert) and nutrient sensing (Fgf21) as well as the knock-out of Akt2, involved in growth signalling and glucose homeostasis. Out of these genetic interventions, FGF-21 overexpression appears to be the most robust since it extends lifespan in both sexes. The other interventions had sex dimorphic effects (Sirt6: male only) or were only tested in one sex (Tert, Akt2).

Based on our initial analysis of GenAge, we performed a literature search for confirmatory studies related to the top genes and pathways identified above. We searched for interventional studies using drugs or viral vectors specifically, because these approaches were not included in GenAge. Only two pathways were supported by such additional evidence, mTOR and telomerase. Somewhat surprisingly, studies targeting the GH/IGF-1 pathway pharmacologically have been less successful, with only one study showing lifespan extension in long-lived mice that was furthermore limited to females (**Duran-Ortiz et al. 2021; Mao et al. 2018**).

We identified studies of the mTOR inhibitor rapamycin based on a recent review (**Selvarani et al. 2021**) and for telomerase activation we searched the literature for published studies. Although the lifespans of most controls were short for both these interventions, comparison with historical controls enabled us to assess their longevity extending properties (**Fig. 5B**). Since a recent meta-analysis reported that metformin fails to extend the lifespan of mice, we used this dataset as a negative control (**Parish and Swindell 2022**). We applied a modified 900-day rule to compare metformin, rapamycin and telomerase activation. 3 out of 9 telomerase studies passed our criteria (38%), 16 out of 30 rapamycin studies (53%) also passed whereas only 1 out of 20 metformin (5%) studies did (**Table S16**).

Finally, comparative analysis of absolute lifespans reveals that drugs do not fully capture the lifespan benefits conveyed by genetic mutations (**Fig. 5A, C** vs **Fig. 5B, D**). In addition, most of these mutations are loss of function rather than gain of function (**Fig. 5D**), suggesting that transgenic mice overexpressing longevity genes are an underexplored area of research.

### Control lifespans over the years – a need for further improvement

Looking at the historical development of mouse lifespan studies, we find that the late 70s and early 80s saw a marked improvement in lifespan (**Fig. 6A**). This is more likely due to improved husbandry rather than a shift towards the use of longer-lived strains since the same trend was observed when we limited our analysis to the popular C57BL/6 strain only (**Fig. 6B**). After this period of marked improvement, lifespan plateaued around 800 days. This increase in lifespan is consistent with a convergence towards a strain-specific optimum. However, we suggest that further improvements in husbandry and mouse lifespan would enable identification of lifespan-extending compounds and interventions with higher confidence and fewer false-positives.

**Figure 6.**
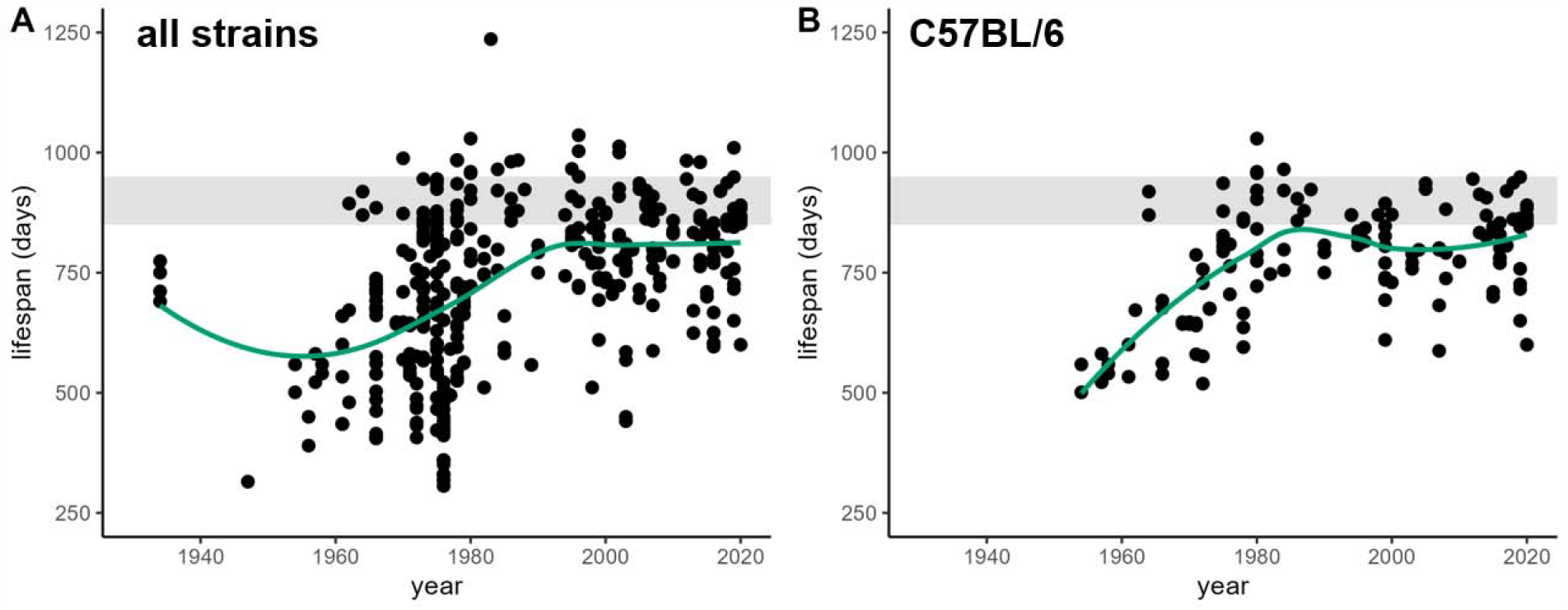
Experimental mouse lifespans improved over time. Reported lifespans in mouse studies improved over the course of the second half of the 20^th^ century. The same trend is seen in an analysis including all mouse strains (A; n=428) and in an analysis limited to studies using C57BL/6 mice (B; n=129). Each datapoint represents the control lifespan of a study or a cohort within a study. Green trend line generated by locally estimated scatterplot smoothing (LOESS) method. The interval between 850 and 950 days is indicated with a shaded area. Data from **Austad (2011), Swindell et al. (2012), Barardo et al. (2017)** and this manuscript.

## Discussion

Although it is conventional wisdom that mouse studies should utilize healthy and long-lived animals, the impact of variation in the lifespan of control animals on experimental outcomes has not been rigorously explored so far. In this work we showed that short-lived controls are prevalent in lifespan studies leading to exaggerated effect sizes of interventions which could affect the reproducibility of these studies.

To evaluate and improve confidence in longevity-extending interventions we propose a 900-day rule for mouse longevity studies. True slowing of aging in mice can only be confidently measured against the backdrop of long-lived controls that are expected to live roughly 900 days (± 50 days), which is the upper end of a healthy normal lifespan. If a study fails the 900-day rule, i.e. an intervention extends the lifespan of a short-lived cohort, we cannot make any claims about aging with confidence except that the tested intervention allowed the animals to reach a lifespan closer to the natural lifespan of a healthy cohort (hence the term longevity-normalizing). In such a case the results have to be interpreted with caution, the study repeated, or the data compared to appropriate historical controls that meet the 900-day rule.

We suggest three explanations for a longevity-normalizing effect. First, the intervention does not affect aging but instead improves the health of animals maintained under sub-optimal conditions, with a genetic predisposition toward short lifespan, or experiencing a diseased state. Second, the intervention has no biological effect and the results are due to regression to the mean or publication bias. Third, the intervention did slow aging, but the effects were overwhelmed by unmeasured factors that lowered the lifespan of both the control and treatment group.

We recognize that no experiment can guarantee, no matter how good the conditions, that the control lifespan will reach close to 900 days. The ITP, for instance, does not always achieve this goal in males. Furthermore, a longevity normalizing effect of an intervention does not preclude it from having health benefits in human populations. It is likely that many people are aging in non-optimal conditions such that longevity-normalizing interventions may have real benefits.

Metformin may be an example of a longevity-normalizing drug, because it works in short-lived mice but not in long-lived mice as shown by application of the 900-day rule. Nevertheless, the drug is associated with numerous health benefits in humans (**Kulkarni et al. 2020**) and we find evidence of synergistic lifespan benefits between rapamycin and metformin in mice.

Our analytical approach produces several other novel insights. We find that many compounds reported to extend mouse lifespan fail to extend lifespan in the ITP upon attempted replication, with the most likely explanation being that the initial results did not pass the 900-day rule. We can also account for many of the sex dimorphic effects seen in the ITP, since males are shorter-lived than females and thus benefit more from longevity-normalizing interventions. Finally, by applying the 900-day rule and comparison with historical controls we were able to identify several promising interventions for further study, e.g. ACE inhibitors, telomerase activation, FGF-21 or rapamycin combinations. Therefore, the use of historical controls is highly recommended especially when the within-study control fails to reach the expected lifespan.

More generally, our approach provides an opportunity to address what is widely appreciated as a “reproducibility problem” in the field. There have been several notable examples where high-profile publications have initially claimed lifespan extension resulting from an intervention only to have subsequent studies fail to reproduce those claims (**Harrison et al. 2021, Strong et al. 2013**). This is particularly problematic in the context of mouse longevity studies, because attempts at replication take several years and require large amounts of resources. Additionally, the intense media and public interest in “anti-aging” regimens means that such reports are often widely disseminated to the general public, often accompanied by direct marketing of products to consumers. Hence, there is an urgent need for clear guidelines to confidently identify lifespan extending compounds.

### Summary and limitations

Although theoretically the reliability of a mouse lifespan study should be proportional to the lifespan of the controls across the whole range of values, we nevertheless see certain advantages in the 900-day rule for practical purposes.

Specifically, the advantages of a simple, binary rule are ease of use and ease of adoption. These often outweigh the disadvantages like lack of precision and explanatory power. One example where this choice was made by convention would be the famous p-value cut-off α=0.05. Such rules should not discourage subject experts from a more thorough exploration of the raw data, while opening the field to a wider number of scientists and audiences.

## Methods

### Data collection and pre-processing

We collected median lifespans from the literature when possible, or mean lifespans when only these were provided by the authors. If neither was provided, we determined median LS from survival curves. Measures of maximum lifespan or mortality doubling time were not considered due to higher statistical uncertainty associated with these. When up-to-date data was not available, as was the case for recent studies of CR and telomerase activation, we performed a systematic literature search to identify studies and extend existing datasets.

All datasets used in this manuscript are summarized in **Table S1** and **Table S2**. Correlation analysis was performed on the level of individual studies or cohorts, not individual animals. We removed datapoints deemed to be of low quality (e.g. no adequate information on strain and sex given). We further cleaned up some datasets as needed, e.g. removing duplicates, or entries with missing references. Furthermore, we excluded the ITP and rapamycin data from DrugAge, which we analyze in more detail elsewhere. For GenAge, whenever multiple cohorts were reported in a paper, we chose the cohort with the highest lifespan for our analysis.

### Analysis, linear regression and outlier removal

We performed Pearson correlation in this study, although the results were comparable using Spearman correlation (**Table S3**). For the ITP dataset, we calculated a p-value using the lmerTest package in R to construct a linear mixed effects model with a random term accounting for cohort year and test center.

To minimize denominator bias, we plot control lifespan against absolute change in lifespan rather than relative change (lifespan^treated^/lifespan^control^), although data is comparable for both (**Table S1**). Outlier removal in **Fig. 2** was performed and R-values are the worst case of leave-one-out analysis.

### Analysis of sex and drug effects in the Interventions Testing Program

Raw data was obtained from the study authors. For the comparison of sex dimorphic effects only treatments that were tested in both sexes were included and the sex-specific survival advantage was calculated as absolute lifespan extension^male-female^. To obtain results unbiased by multiple testing of one and the same drug, we randomly chose a lifespan study within each drug class for our analysis.

### Resampling to model regression to the mean

Whenever the control group is longer-lived than the true population mean by chance, the treatment group will be on average closer to the mean and thus shorter-lived. The inverse will apply to short-lived controls giving rise to a negative correlation between control group lifespan and lifespan extension of the treated group (regression to the mean). To compare the observed lifespan data with a theoretical null distribution showing such regression to the mean effects, we performed a bootstrap analysis. Given the underlying lifespan distribution of the control cohort, we resampled from this control population with replacement and group sizes matching the actual experiment. The effect of regression to the mean is then estimated by comparing the slope of the resampled regression line with the slope of the observed regression line. To this end, we calculated a z-score for the difference between the slopes and used this to compute a two-tailed p-value.

### Defining lifespan gold standards

An idealized “healthy lifespan” of a mouse is defined as the longest median lifespan that a cohort of lean animals can achieve without slowing the rate of aging per se. Although this quantity is not knowable, we can gain an intuition by studying historical lifespan data. It is likely that a healthy cohort asymptotically converges towards a species- and strain-specific median lifespan optimum. Indeed, improvements in general health and husbandry lead to rectangularization of the survival curves and convergence towards this optimum in both mouse experiments (Hayflick and Finch 1977) and human populations (**Yashin et al. 2012, Myers et al. 1984**).

## Abbreviations

ITP: Interventions Testing Program
CR: caloric restriction
ILSXISS: recombinant inbred cross of ILS (Inbred Long Sleep, ILS) and ISS (Inbred Short Sleep, ISS) mice
FGF-21: fibroblast growth factor 21
GH: growth hormone

## Acknowledgments

We thank VitaDAO for financial support, Giuliani Alessandro, David B. Allison, anonymous reviewers and Michael Rae for constructive feedback. We also thank Rich Miller, Basten Snoek, Arlan Richardson and Daniel Promislow for providing the raw lifespan data.

