## Supplemental Data for "The impact of short-lived controls on the interpretation of lifespan experiments and progress in geroscience"

**Supplementary figures**

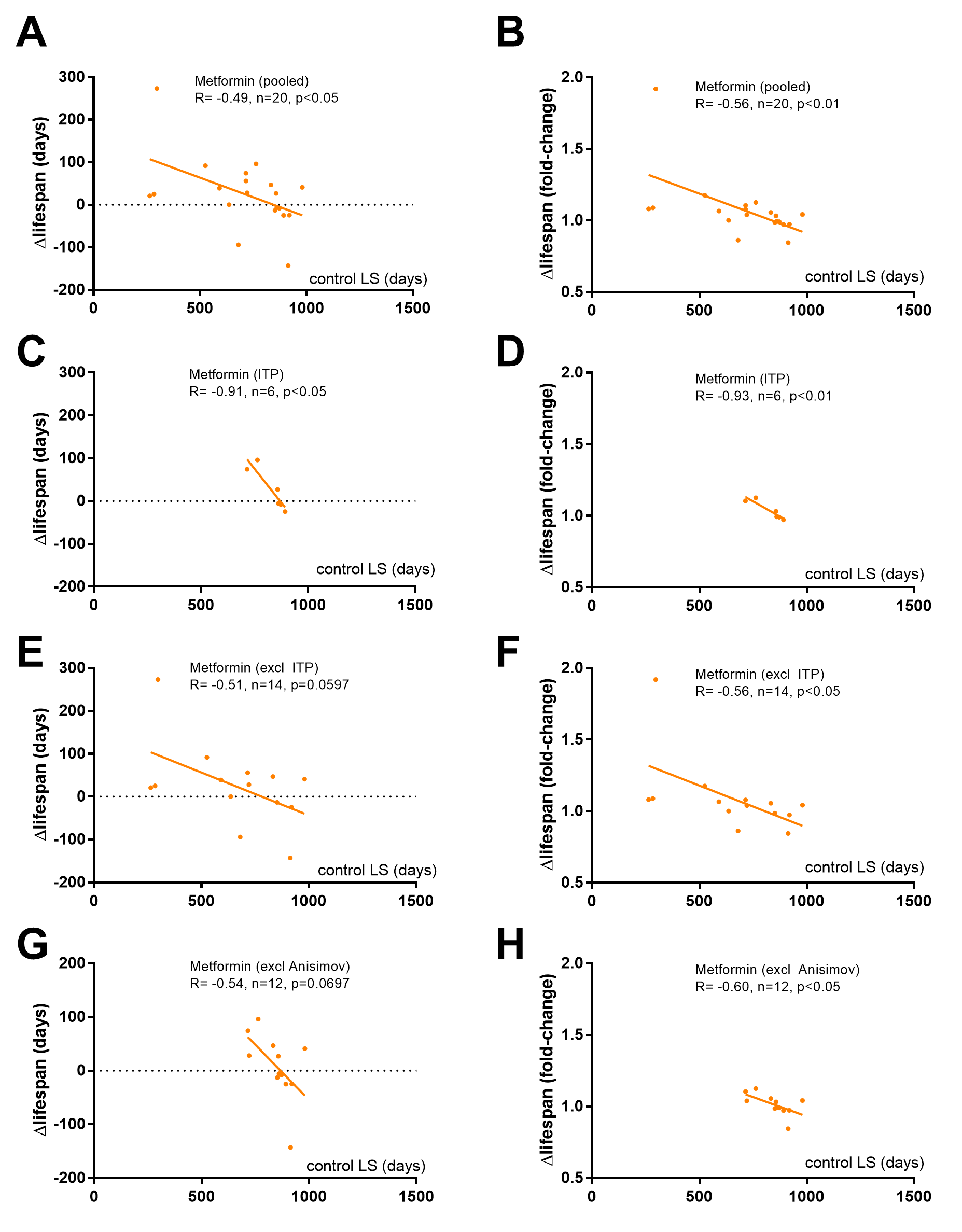

**Fig. S1. Metformin fails to extend the lifespan of long-lived mice**

In this figure we plot the lifespan of control mice on the X-axis against the relative or absolute change in lifespan with metformin treatment on the Y-axis. In all cases, the effects of metformin are highly dependent on the lifespan of control mice and the drug has no benefit in long-lived animals.
(A, B) The relative (A) and absolute (B) change in lifespan with metformin treatment is plotted (n=20).
(C, D) The relative (C) and absolute (D) change in lifespan with metformin treatment is plotted. Data based on our analysis of the ITP (n=6).
(E, F) The relative (E) and absolute (F) change in lifespan with metformin treatment is plotted after excluding the ITP from the analysis (n=14).
(G, H) The relative (G) and absolute (H) change in lifespan with metformin treatment is plotted after excluding the data by Anisimov et al. from the analysis (n=12).

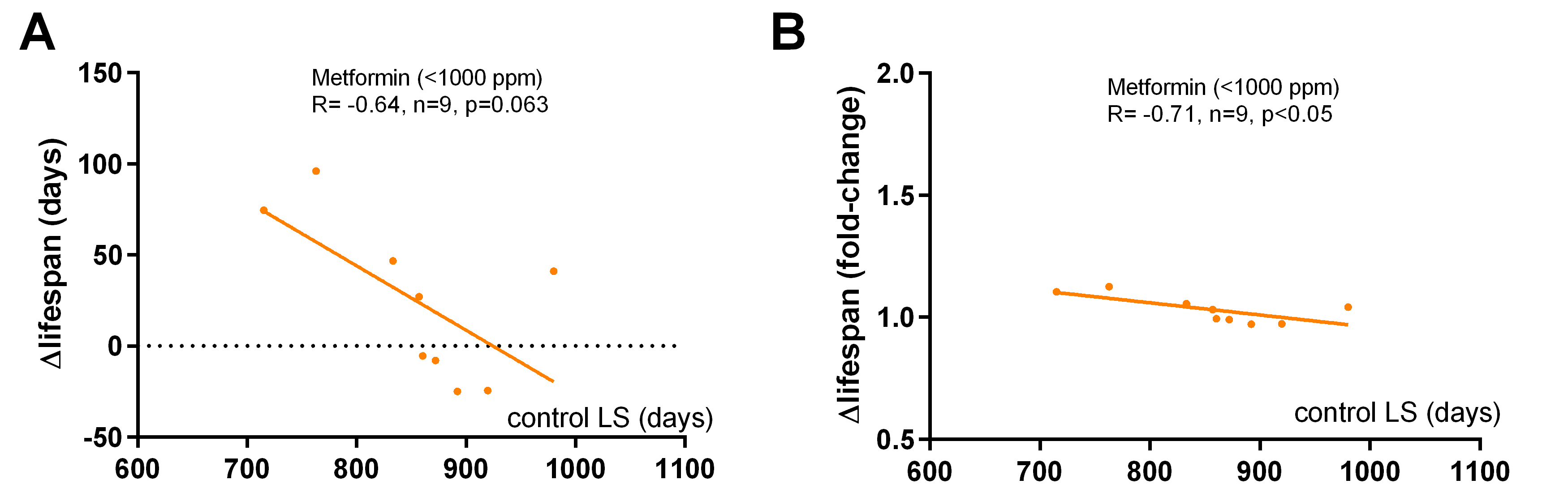

**Fig. S2. Low doses of metformin fail to extend the lifespan of long-lived mice**
In this figure we plot the lifespan of control mice on the X-axis against the absolute (A) or relative (B) change in lifespan with metformin treatment (<1000 ppm) on the Y-axis. The effects of metformin are highly dependent on the lifespan of control mice and the drug has no benefit in long-lived animals (n=9).

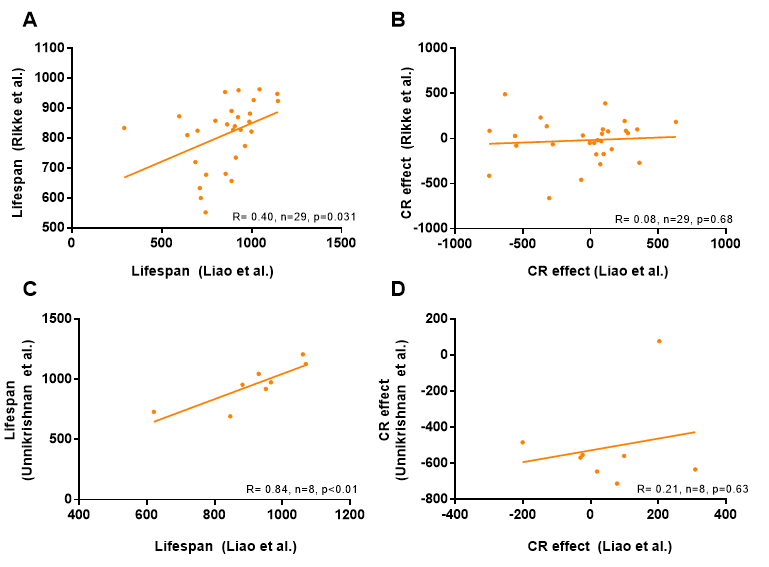

**Fig. S3. Control lifespans are similar for ILSXISS strains tested in different studies, whereas the response to caloric restriction is not**
The lifespans of control ILSXISS mice are comparable between Rikke et al. and Liao et al. (A) or between Liao et al. and Unnikrishnan et al. (C). In contrast the response to CR (lifespan shortening or extension in days) is not comparable between Rikke et al. and Liao et al. (B) or between Liao et al. and Unnikrishnan et al. (D). The correlation between the results of Rikke et al. and Unnikrishnan et al. is not shown because only three strains were evaluated by both studies.

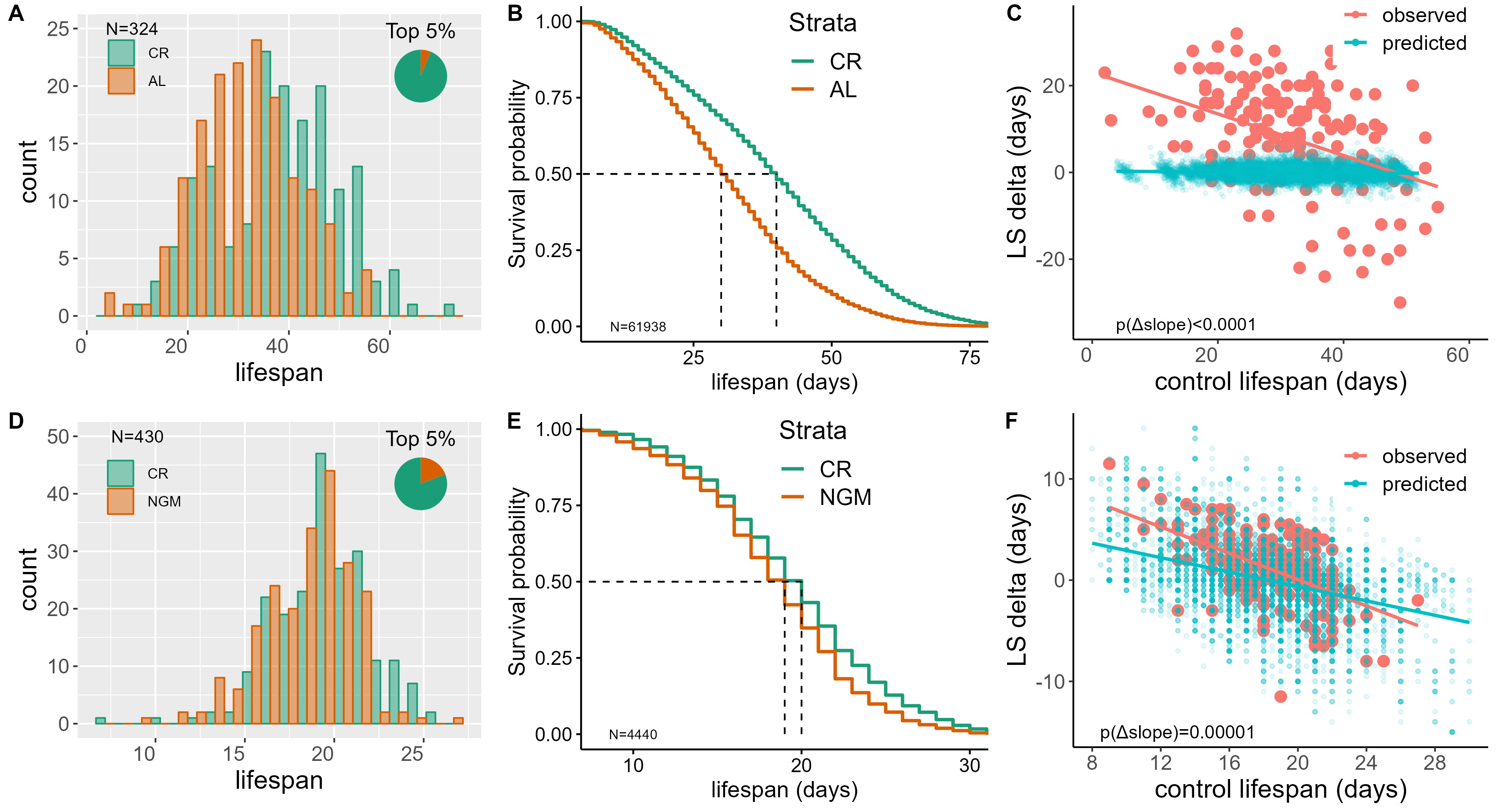

**Fig. S4. Caloric restriction extends the lifespan of flies and worms, with long-lived strains benefiting less**
Caloric restriction (CR) extends the lifespan of flies and most of the longest-lived cohorts in the study of Jin et al. (2020) are from the CR group (A, n=324 cohorts). When pooling the individual level data, we also see a significant lifespan extension under CR (B). Longer-lived flies responded less favorably to CR than expected (C). Similarly, CR extends the lifespan of worms and most of the longest-lived cohorts in the study of Snoek et al. (2019) are from the CR group (D, n=430 cohorts) and the same is true when pooling data from all the individual worms (E). As with flies, longer-lived worms responded less favorably to CR than expected (F). To test whether regression to the mean can explain exaggerated benefits in short-lived strains we resampled quasi-lifespan experiments from the control population. The resampled synthetic data (blue) is shown for flies (C) and worms (F) with the observed datapoints overlaid (pink).

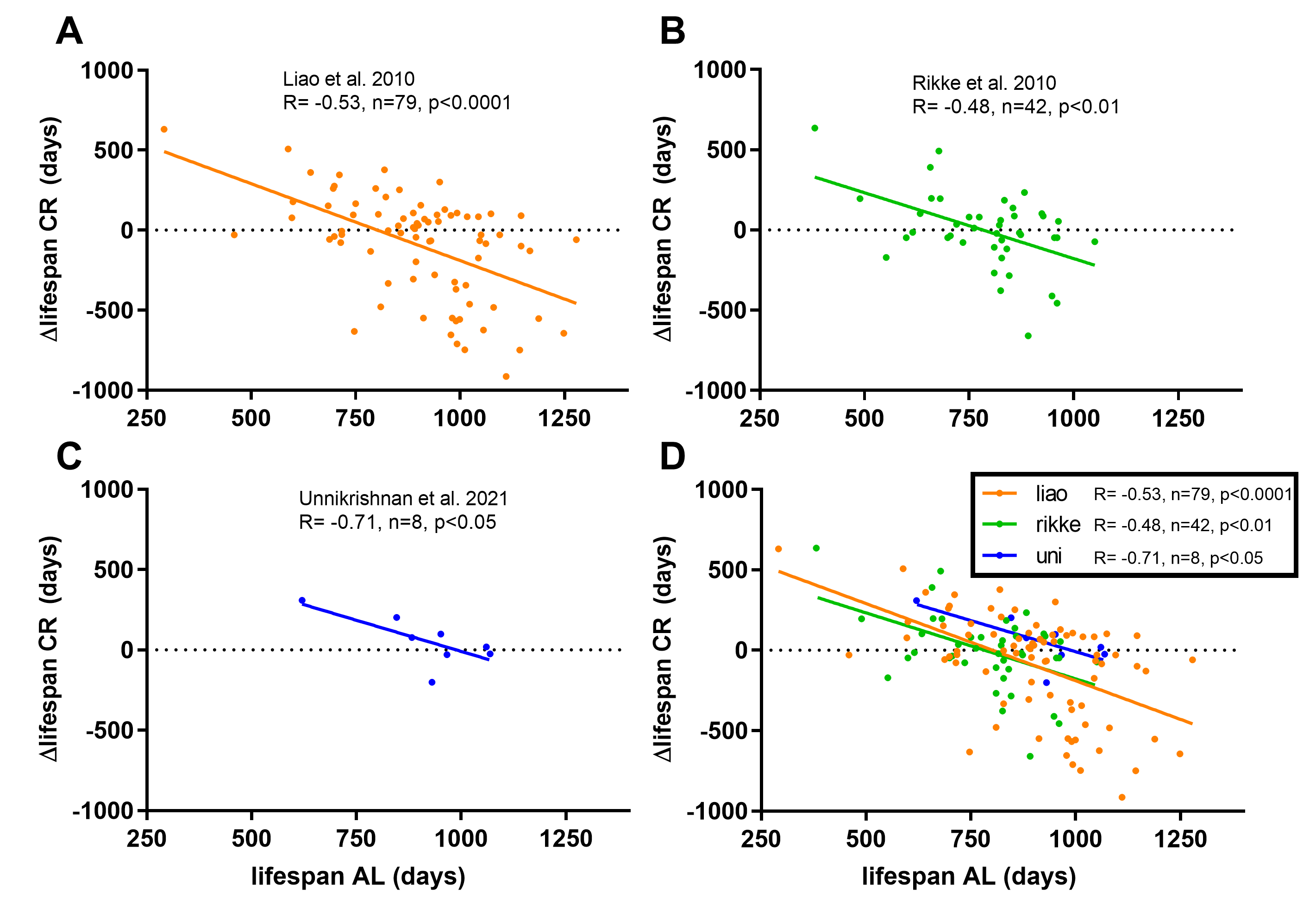

**Fig. S5. Ad libitum lifespans of ILSXISS mice predict the response to caloric restriction in three different datasets**
In this figure we plot the lifespan of control mice fed ad libitum (AL) on the X-axis against the absolute change in lifespan with caloric restriction (CR) on the Y-axis (Δlifespan). Each point represents a particular ILSXISS strain. In all three studies long-lived strains respond less favorably to CR.
(A) Based on a re-analysis of mouse data from **Liao et al. (2010)**.
(B) Based on a re-analysis of mouse data from **Rikke et al. (2010)**.
(C) Based on a re-analysis of mouse data from **Unnikrishnan et al. (2021)**.
(D) All three datasets (A-C) superimposed.

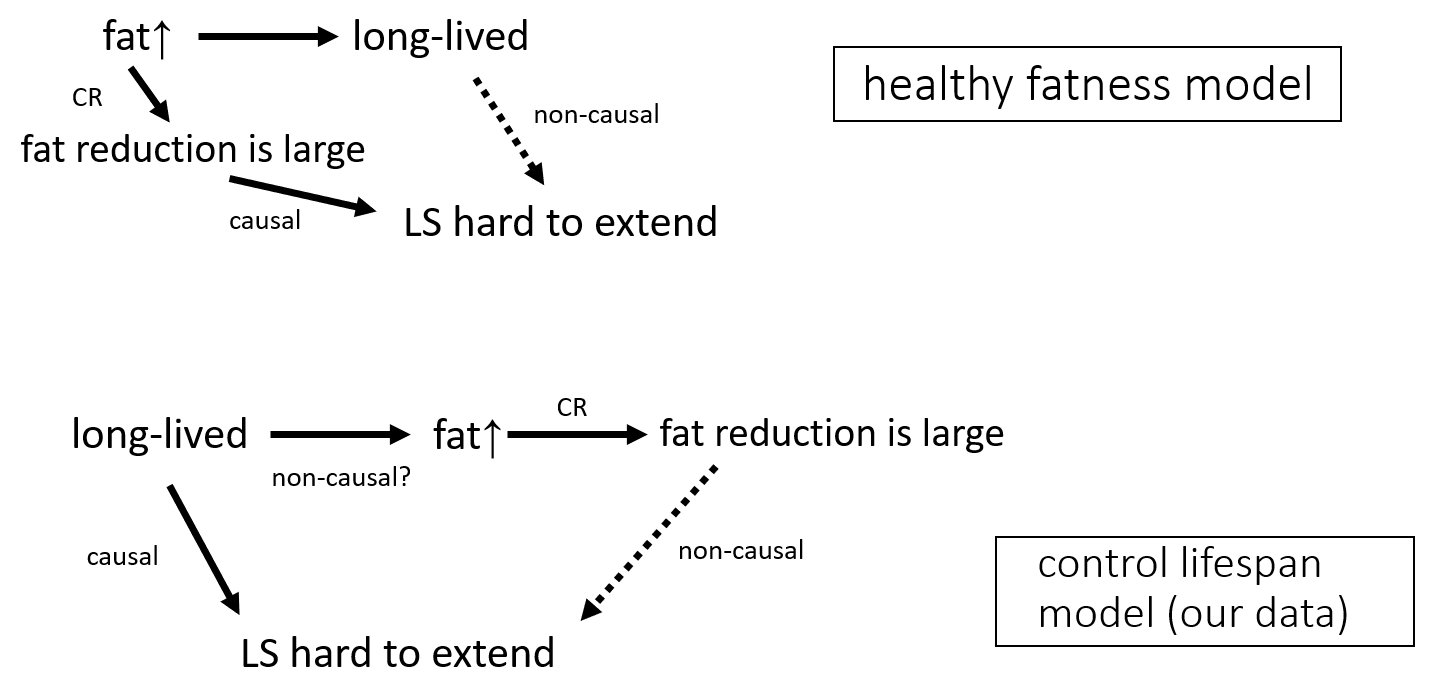

**Fig. S6. Two models that can account for the ILSXISS data: healthy fatness and control lifespans**
The inability of CR to extend the lifespan in certain ILSXISS strains is most likely explained through mix of two different models.
(A) Original healthy fatness hypothesis as raised **Liao et al. (2011)**. A high amount of body fat is beneficial to longevity, therefore a large reduction in body fat under CR is harmful. There is no link between longevity per se (control group lifespan) and failure to further extend lifespan under CR in this model.
(B) In our model, the lifespan of mice that are already long-lived is difficult to extend. Long-lived ILSXISS mice happen to be fat – whether this is causal or purely coincidental – and since adipose tissues is preferably lost during CR this will create the illusion that a loss in adipose tissue is harmful.

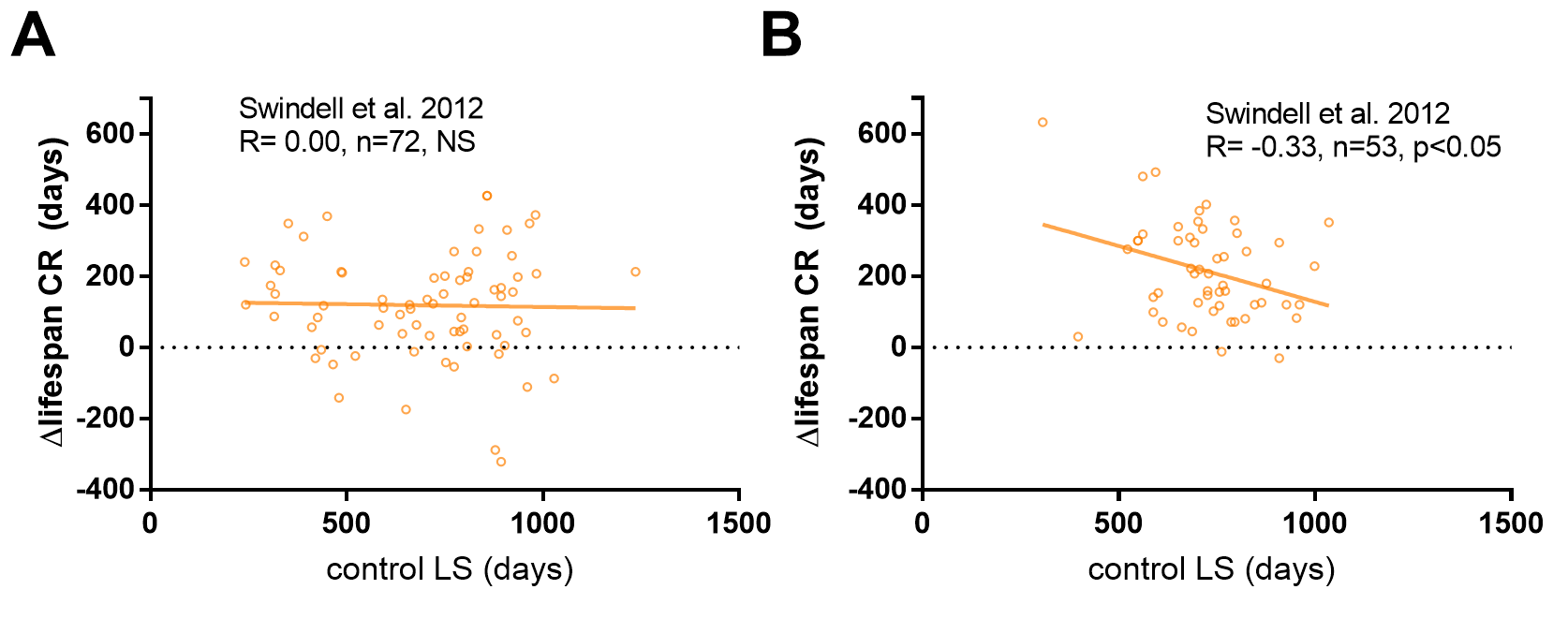

**Fig. S7. Limited association between control lifespans and the effect of caloric restriction or drug interventions in meta-analyses**
In this figure we plot the lifespan of control mice (AL) on the X-axis against the absolute change in lifespan with caloric restriction or drug interventions on the Y-axis (Δlifespan). Control lifespans have a significant effect on lifespan extension under CR in rats (A) but not mice (B).
(A) Based on a re-analysis of mouse data from **Swindell et al. (2012)**.
(B) Based on a re-analysis of rat data from **Swindell et al. (2012)**.

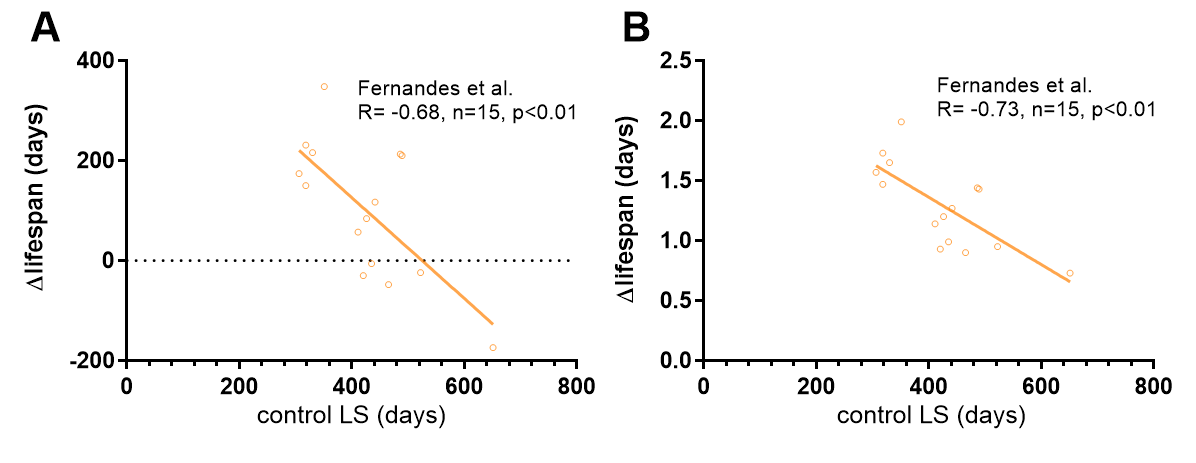

**Fig. S8. Short-lived mice show an exaggerated benefit from caloric restriction**
In this figure we plot the lifespan of ad libitum fed control mice on the X-axis against the relative (A) or absolute (B) change in lifespan with caloric restriction on the Y-axis (Δlifespan). Shorter-lived B/W and DBA/2f mice benefited more from caloric restriction. Data based on several studies published by Fernandes et al. (1976).

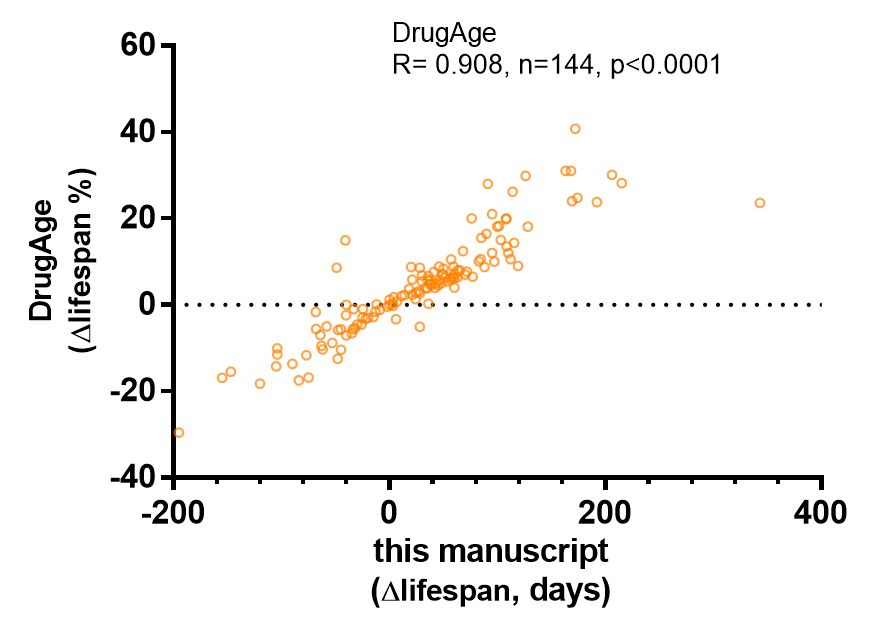

**Fig. S9. Relative and absolute changes in lifespan are correlated when extracted by two different methods (DrugAge data)**
Relative mouse lifespan extension reported in DrugAge (y-axis) and absolute lifespan changes reported in this paper (x-axis). The above data is based on a cleaned-up version of DrugAge excluding rapamycin and ITP data. N=144. Even though we used different methods to extract lifespan data from the DrugAge studies there was nevertheless good agreement between our approach (absolute change) and the original DrugAge database (reporting relative change).

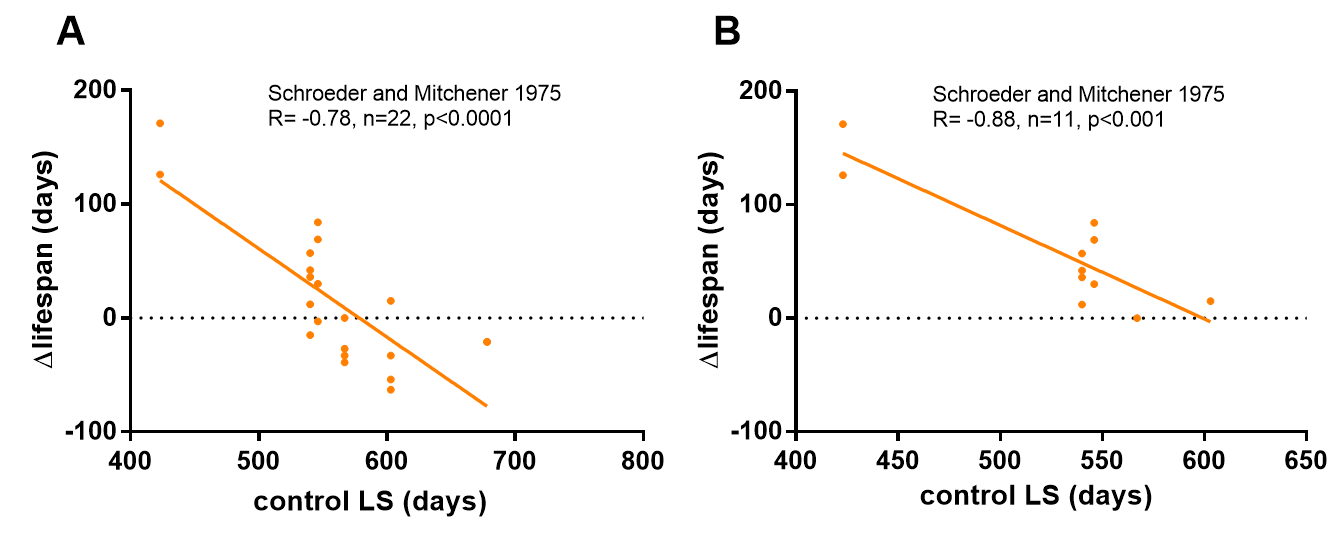
**Fig. S10. Short-lived Swiss mice show an exaggerated benefit from the administration of metals in the diet**
In this figure we plot the lifespan (LS) of control mice on the X-axis against the absolute change in lifespan with metal administration on the Y-axis.
A) Correlation between control lifespan of Swiss mice and ΔLS after treatment with different metals (n=22) in the study of Schroeder and Mitchener 1975.
B) Same as in (A) but with lifespan-shortening treatments excluded.

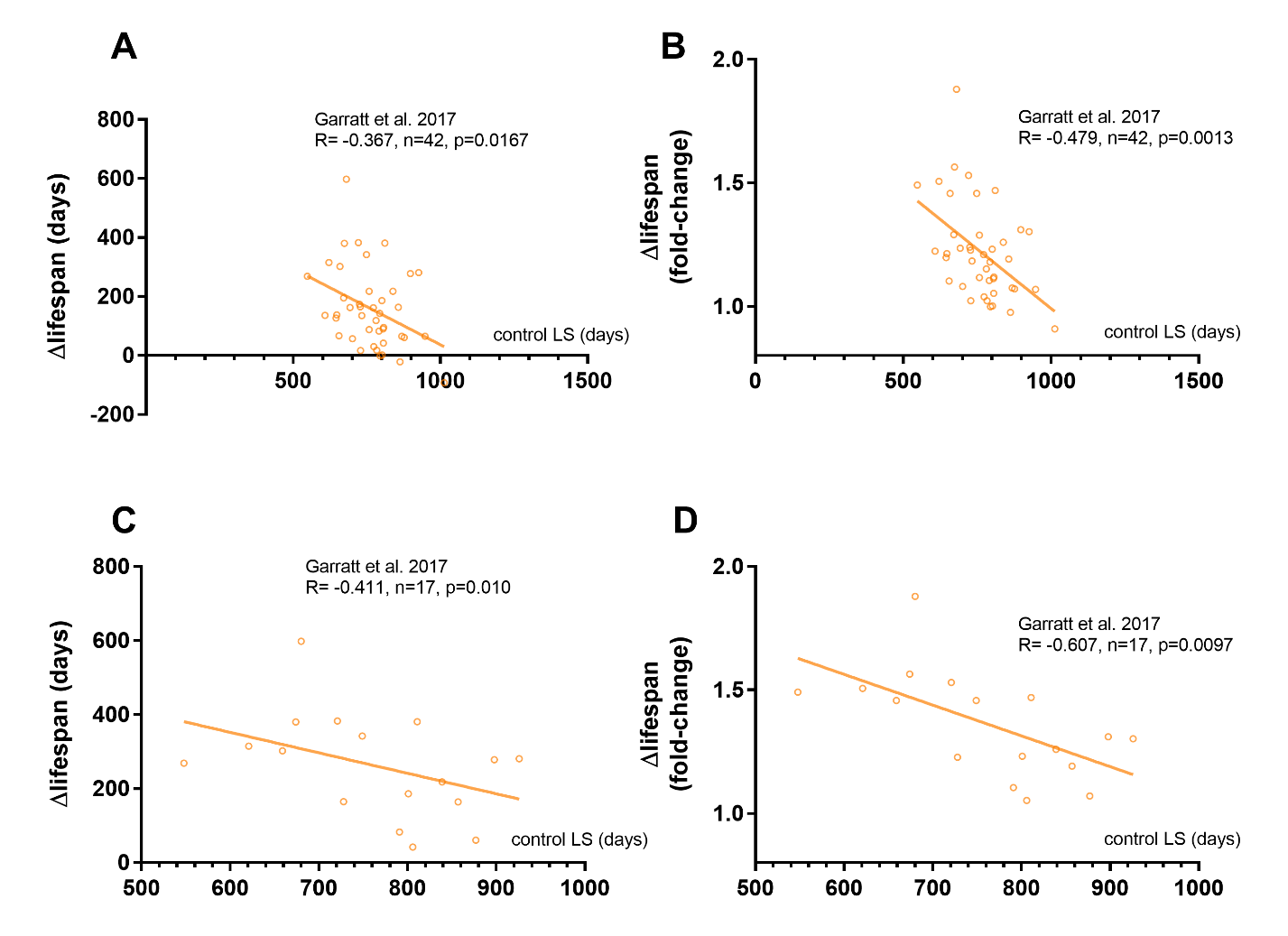
**Fig. S11. Control lifespans predict lifespan extension of growth hormone and IGF-1/IRS mutants**
In this figure we plot the lifespan (LS) of control mice on the X-axis against the absolute change in lifespan with genetic mutations of the somatotropic axis on the Y-axis (Δlifespan). Each point represents a cohort of mice. The control lifespan predicts the response to mutations in the IGF-1/IRS pathway (A, B) and in the GH pathway (C, D). Data expressed as absolute change (A, C) or relative change (B, D).

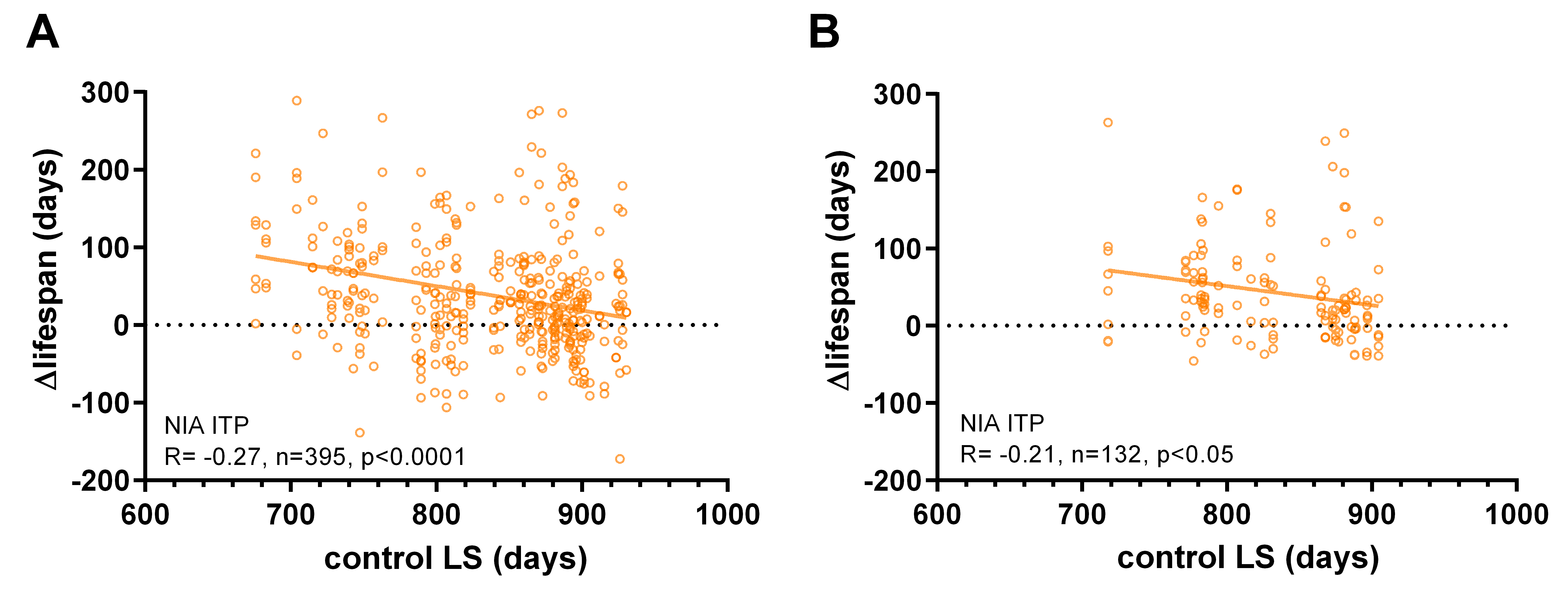

**Fig. S12. The impact of control lifespan variation is diminished in pooled data from the Interventions Testing Program**
We plot the lifespan of mice in the Interventions Testing Program (ITP) study on the X-axis (orange dots) against the change in lifespan with drug treatments on the Y-axis (Δlifespan in days). The negative correlation between control lifespan and treatment outcome is more robust in an analysis that treats each testing site as an independent datapoint (A) rather than pooling lifespans from all testing site (B).

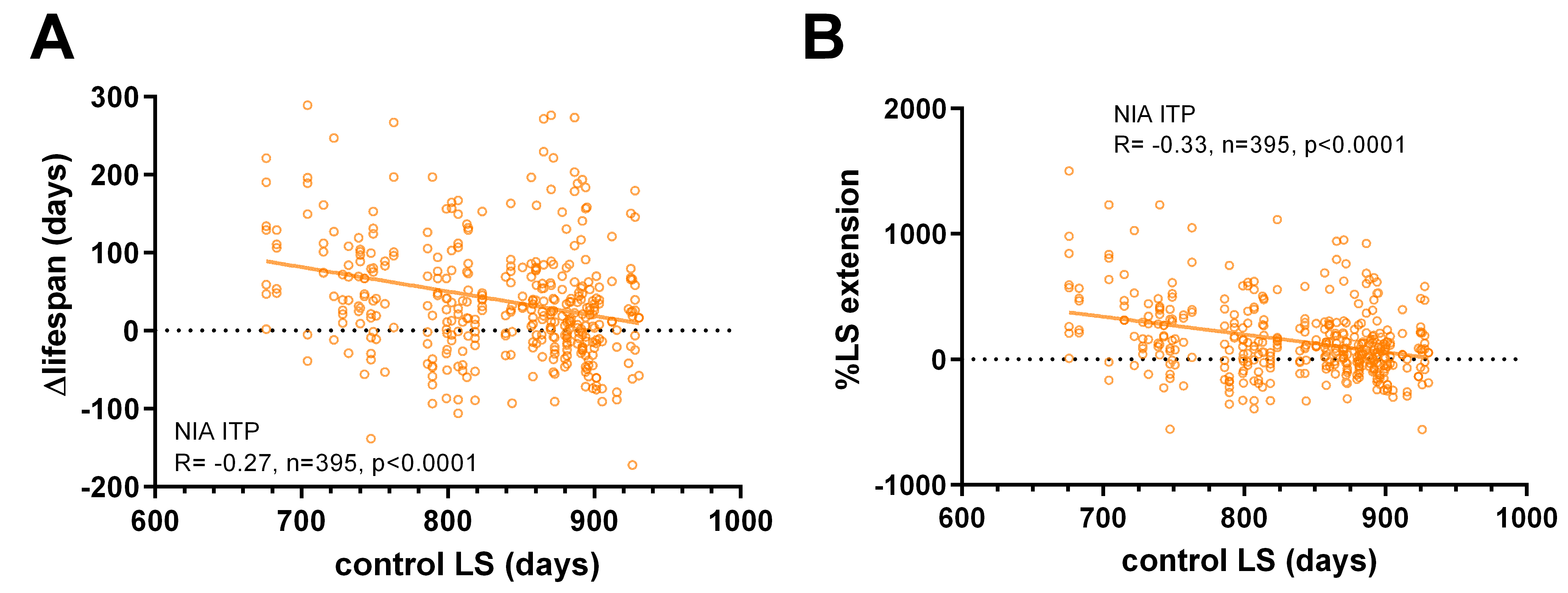
 **Fig. S13. Longer-lived cohorts of UM-HET3 mice show less pronounced lifespan extension in the Interventions Testing Program**
We plot the lifespan of mice in the Interventions Testing Program (ITP) study on the X-axis (orange dots) against the change in lifespan with drug treatments on the Y-axis (Δlifespan in days). Each point represents a unique combination of drug x gender x testing site. The negative correlation between control lifespan and treatment outcome is seen when lifespan extension is plotted as absolute (A) or relative (B) change. The data in Fig. 13A is equivalent to the data in Fig. 12A.

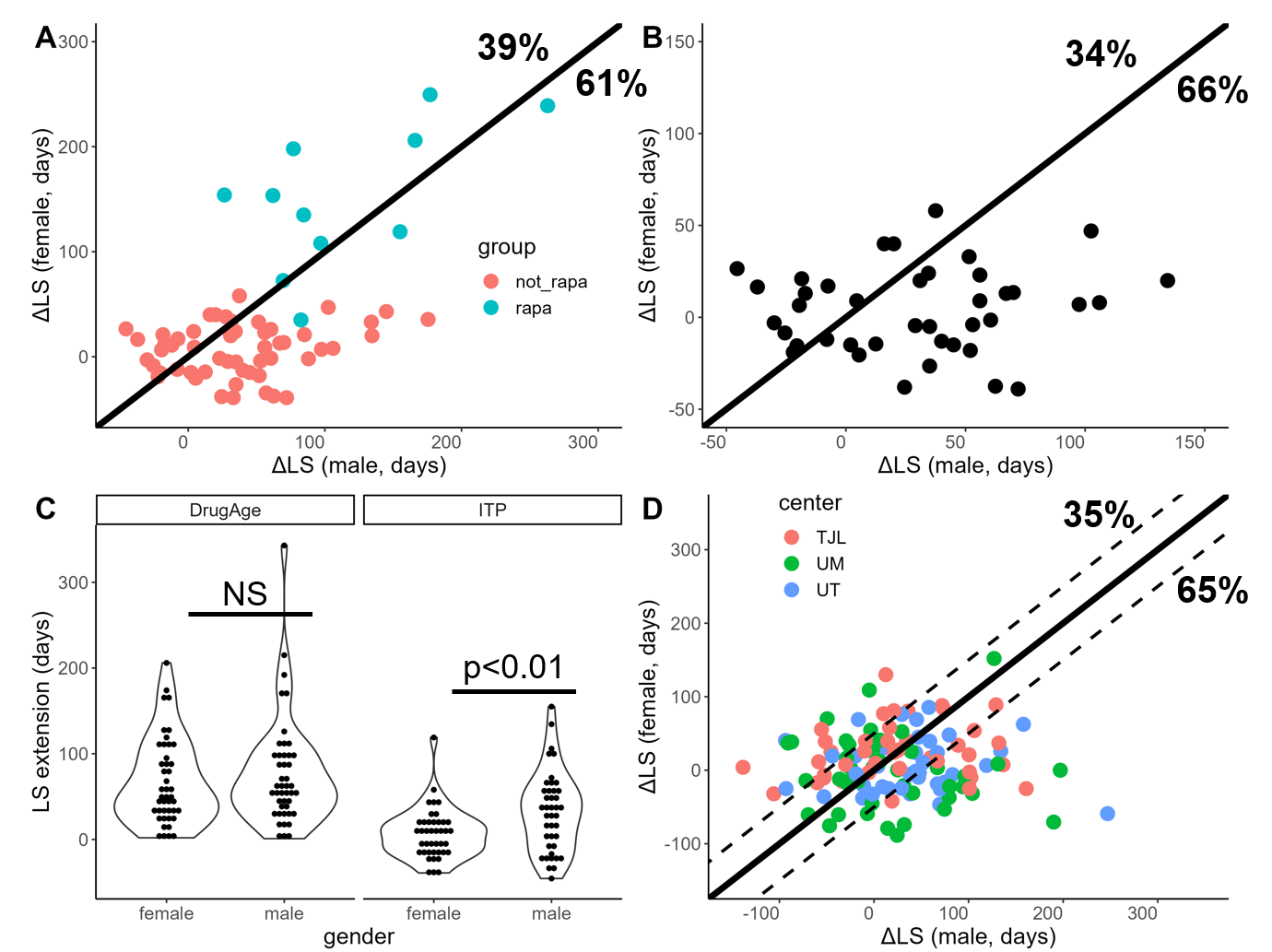

**Fig. S14. Male mice in the Interventions Testing Program benefit more from interventions than female mice**
Here we plot the absolute change in lifespan (LS) with different treatments for male mice and female mice. Each dot represents the results from a mouse cohort. Male mice generally respond better to treatments than do female mice.
A) Treatments pooled across sites (n=64). Cohorts treated with rapamycin are colored in turquoise (“rapa”) and the remaining cohorts in red (“not_rapa”).
B) Unique treatments pooled across sites (n=41). If a drug was tested more than once by the Interventions Testing Program (ITP), these redundant treatments were excluded from the analysis.
C) Lifespan extension in the DrugAge cohort (n=58 female and n=81 male cohorts) and in the ITP across genders (n=143 per gender; each site is an independent datapoint).
D) Unique treatments colored by site (n=123). Percentages in the figure refer to the fraction of treatments outside of the dashed line. Among treatments favoring males, most datapoints were from the University of Texas Health Science Center (UT) testing site (42%) and the University of Michigan (UM) testing site (34%), rather than the Jackson Laboratory (TJL) testing site (24%).

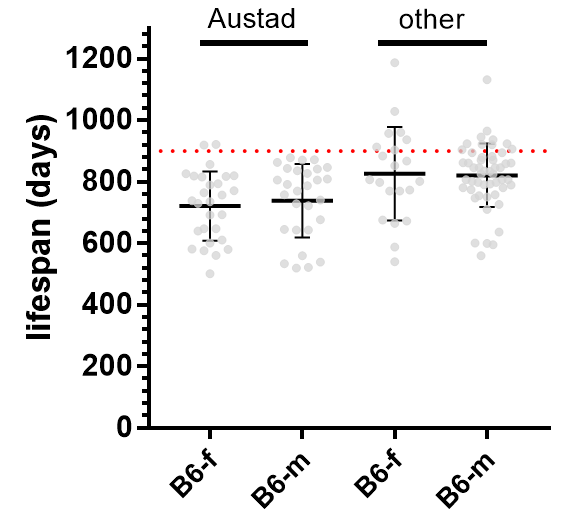

**Fig. S15. Lifespans of** **C57BL/6 mice show no overt sex dimorphism**
Although sexually dimorphic lifespans are observed under some specific conditions and in some datasets, when pooling data from all sources there is no clear sex difference between male (B6-m) and female (B6-f) mice. The “other” group contains pooled data from **Swindell et al. (2012)**, DrugAge (**Barardo et al. 2017**) and from our own analysis. A lifespan of 900 days is indicated with a dashed red line. Errors bars show mean ± SD.

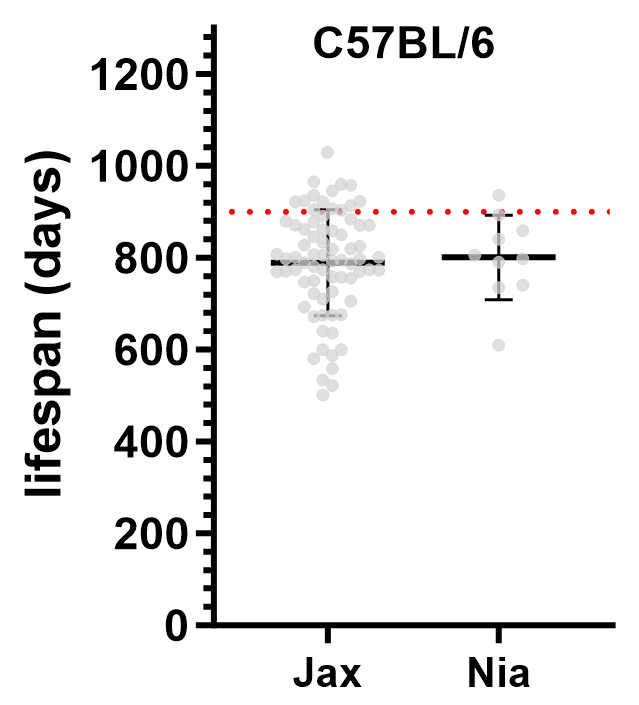

**Fig. S16. Lifespans of C57BL/6J and C57BL/6Nia mice are comparable**
Even though there are some physiologic differences between the Jax and the Nia substrains of C57BL/6 mice, their lifespans are comparable. A lifespan of 900 days is indicated with a dashed red line. Data from **Austad (2011)**, **Swindell et al. (2012)**, DrugAge (**Barardo et al. 2017**) and from our own analysis. Errors bars show mean ± SD.

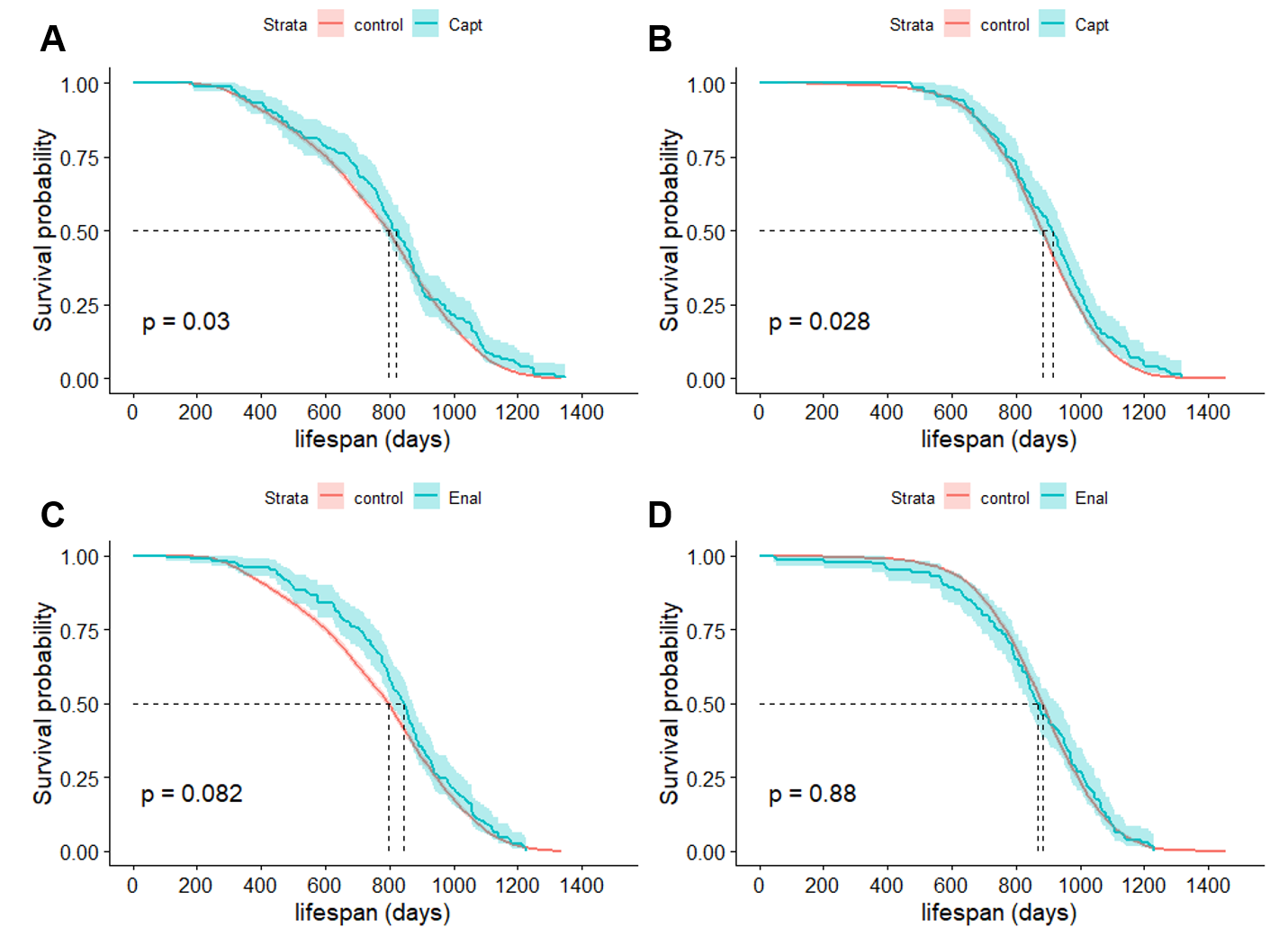

**Fig. S17. Angiotensin-converting enzyme (ACE) inhibitors extend the lifespan of male mice in the Interventions Testing Program when compared to historical controls**
Captopril (Capt) significantly extends the lifespan of male (A) and female (B) mice in the Interventions Testing Program (ITP). Enalapril (Enal) extends the lifespan of male mice (C), although the result does not reach significance, and does not affect the lifespan of female mice (D). P-value by log-rank test between drug and all pooled historical controls from the ITP. 95% confidence interval plotted for both drugs and controls.

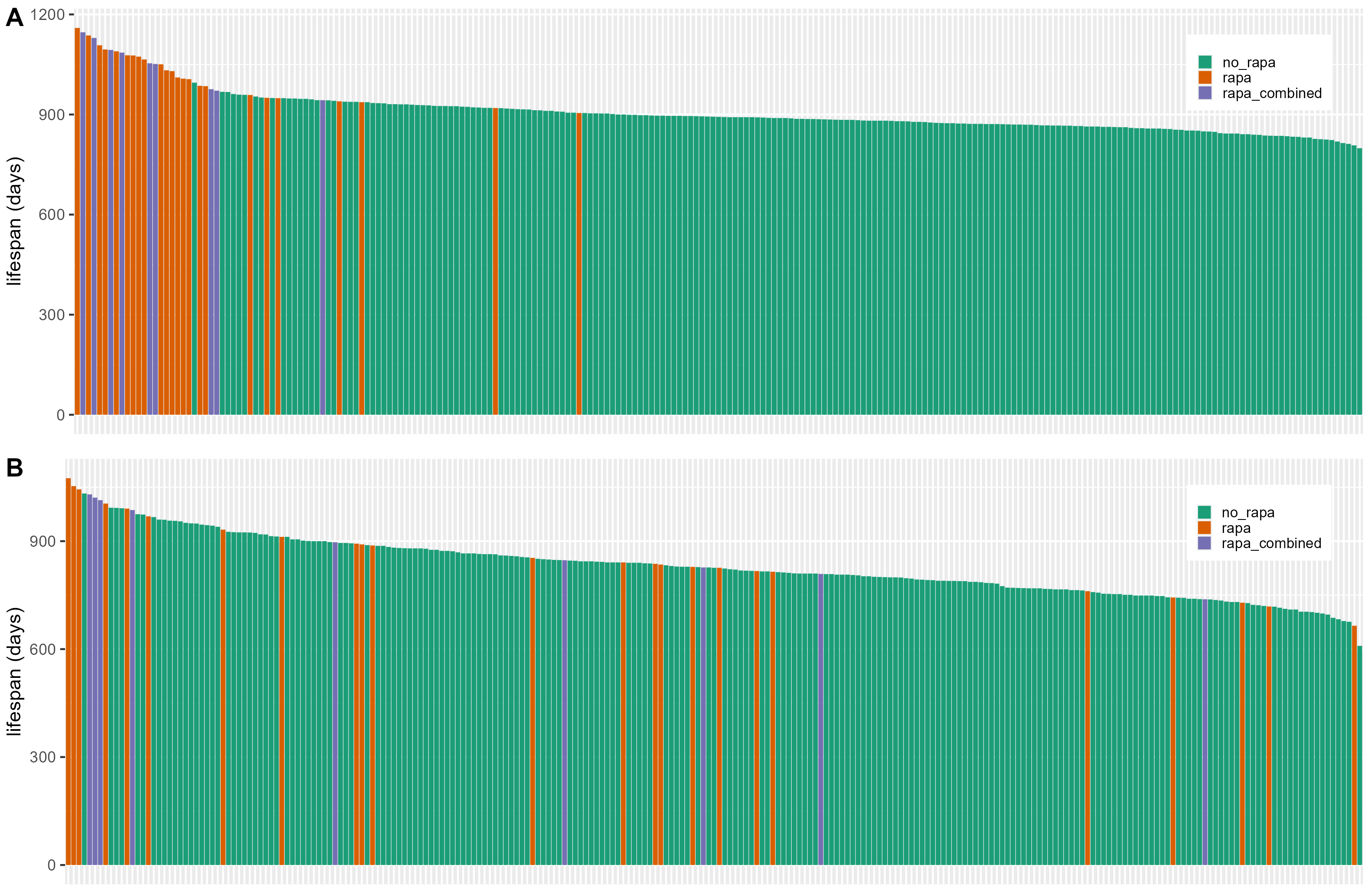

**Fig. S18. Rapamycin combination therapies lead to higher lifespan extension in the ITP than does rapamycin alone**
Rapamycin combination therapies (“rapa_combined”) lead to higher lifespans in female UM-HET3 mice (A) than does rapamycin (“rapa”) by itself (p=0.076 for difference in rank), while this advantage is less pronounced in male mice (p>0.10, B). n>230, compounds ordered by the lifespan of the treated group.a

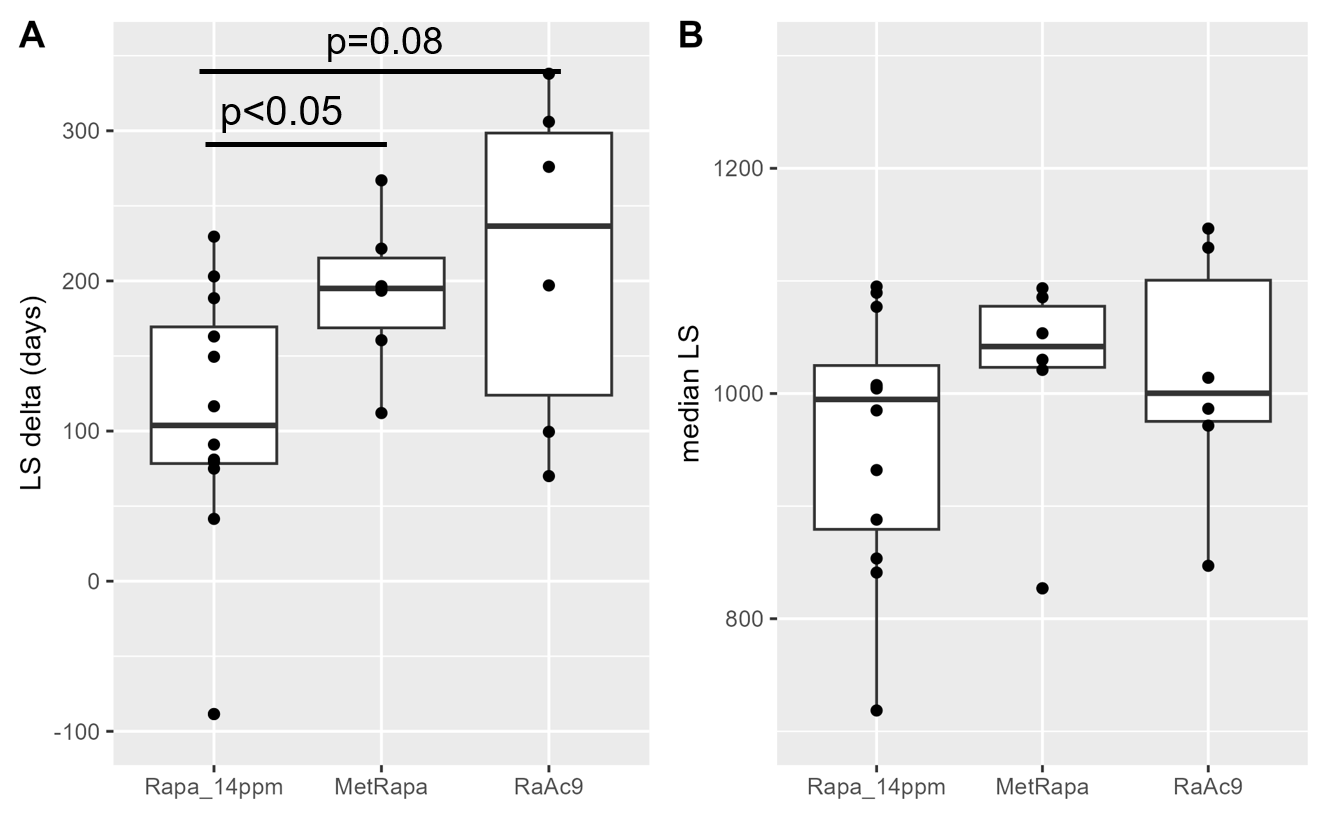

**Fig. S19. Combination therapies lead to higher lifespan extension in the ITP**
Combination therapies lead to numerically higher lifespan extension (A) and absolute lifespans than do single treatments (B). In the combined metformin and rapamycin group (MetRapa) rapamycin was given to UM-HET3 mice at 14 ppm and metformin at 1000 ppm. In the combined rapamycin and acarbose group (RaAc9) rapamycin was given at 14 ppm and acarbose at 1000 ppm. Data for the 14 ppm rapamycin dose was pooled from two cohorts (C2006, C2009). All treatments were started at 9 months of age in both genders. p>0.10 unless indicated.

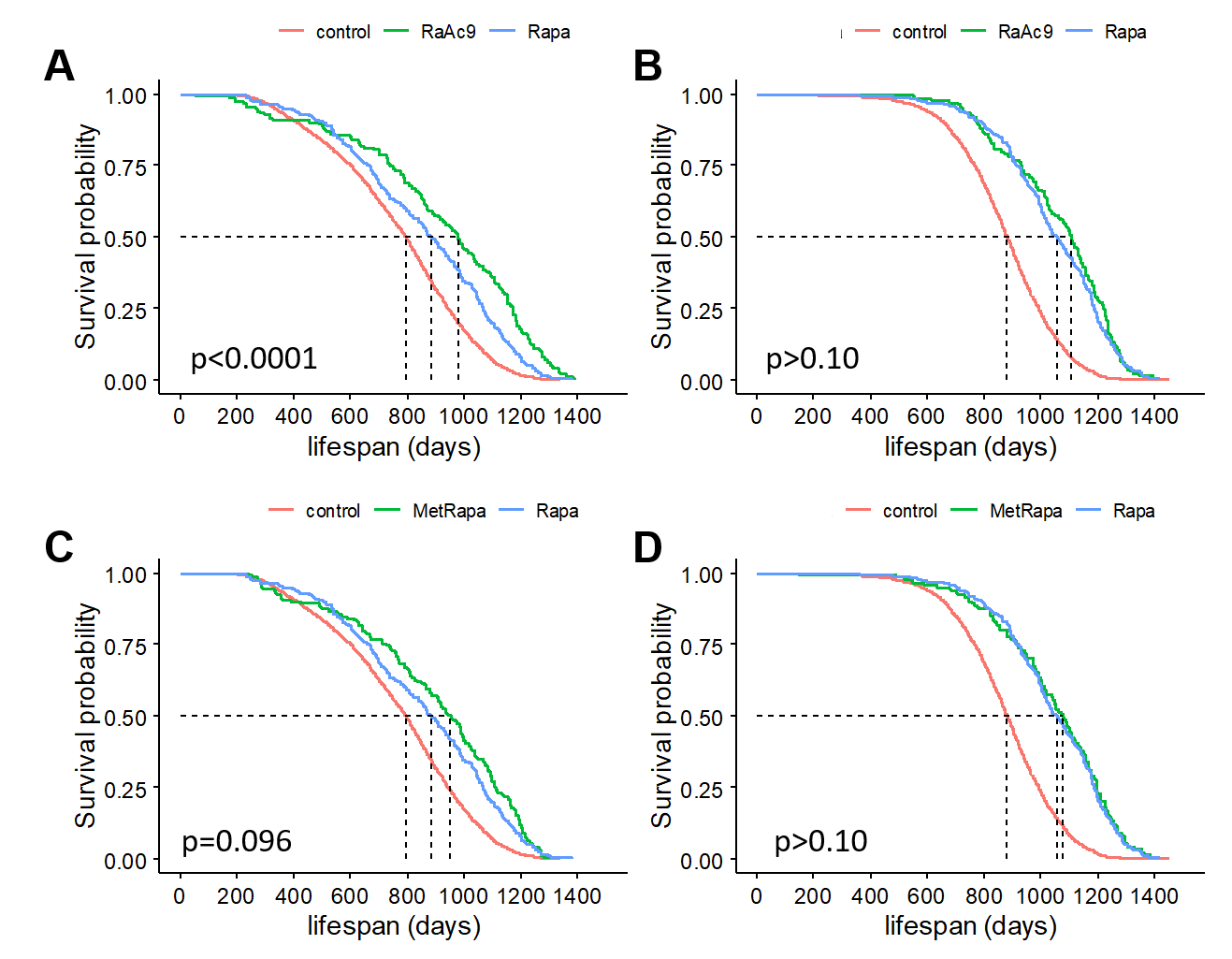

**Fig. S20. Rapamycin combination therapies extend the lifespan of male mice in the Interventions Testing Program when compared to rapamycin alone**
The combination of rapamycin and acarbose significantly extends the lifespan of male (A) and female (B) mice as compared to rapamycin alone. Rapamycin combined with metformin extends the lifespan of male mice (C) as compared to rapamycin alone, although the result does not reach significance. Lifespan of female mice is not further improved by the combination (D). P-value by log-rank test between rapamycin and rapamycin combination treatment. Pooled historical controls from the Interventions Testing Program (ITP) are shown for comparison (red).

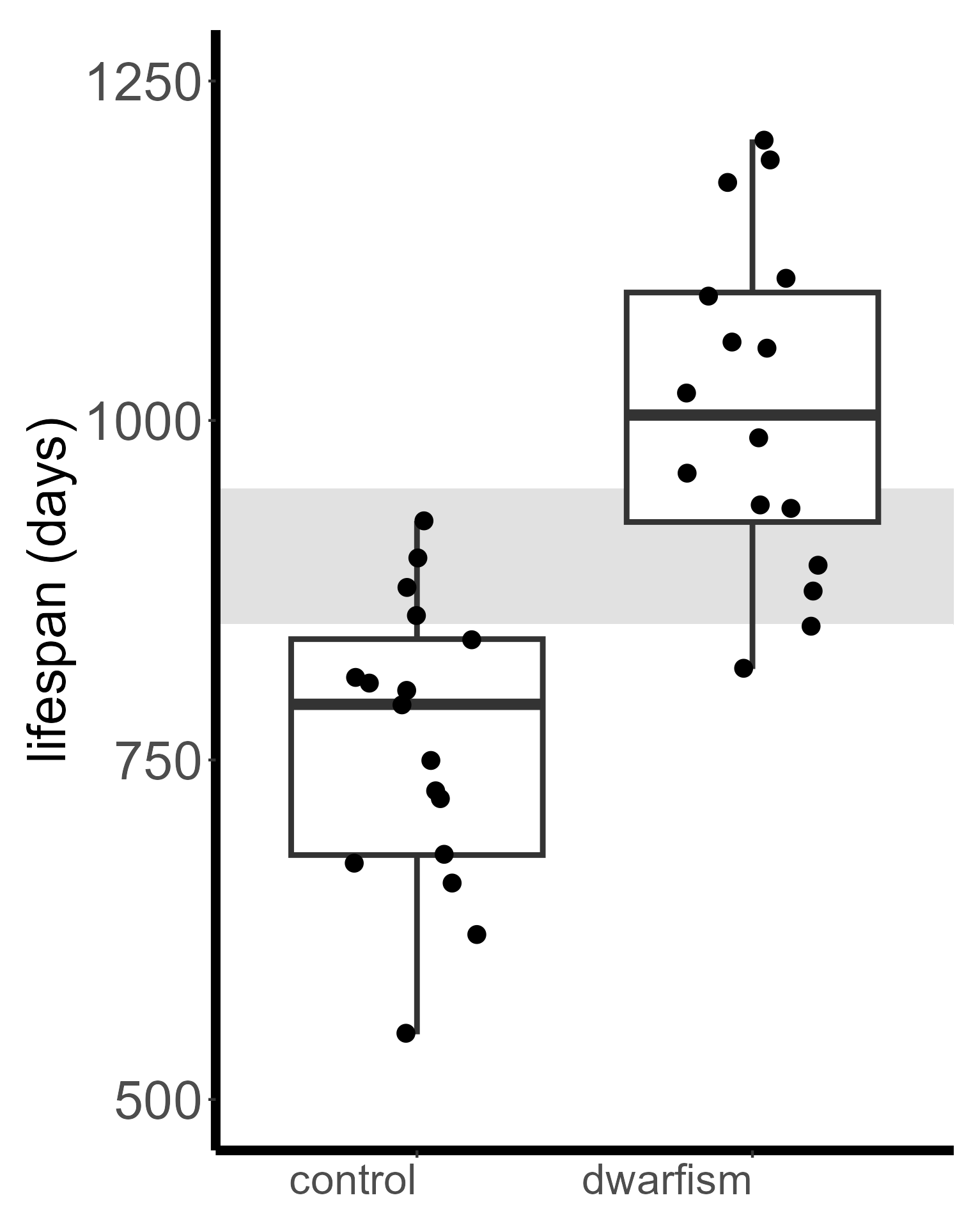
 **Fig. S21. Pituitary dwarfs are unusually long-lived compared to historical controls**
We replot data from the meta-analysis Garratt et al. (2017), showing here that lifespans of pituitary dwarfs, including Ames, Snell and GHRKO dwarfs, are often above 950 days beating the benchmark for historical controls.

| **species** | **strains (dataset)** | **reference** | **abs** | **rel** | **N** |
| --- | --- | --- | --- | --- | --- |
| mice | metformin | Parish and Swindell 2022 | -0.49 | -0.56 | 20 |
| yeast | deletion mutants | Schleit et al. 2013 | -0.24 | -0.19 | 167 |
| flies | DGRP inbred panel | Jin et al. 2020 | -0.38 | -0.57 | 161 |
| Worms | inbred panel | Snoek et al. 2019 | -0.55 | -0.62 | 215 |
| mice | ILSXISS inbred panel | Liao et al. 2010 | -0.58 | -0.53 | 79 |
| mice | ILSXISS inbred panel | Rikke et al. 2010 | -0.56 | -0.48 | 42 |
| mice | ILSXISS inbred panel | Unnikrishnan et al. 2021 | -0.71 | -0.80 | 8 |
| mice | Swiss mice | Schroeder et al. 1975 | -0.78 | -0.80 | 22 |
| mice | B/W and DBA/2f mice | Fernandes et al. 1976 | -0.68 | -0.73 | 15 |
| mice | various (DrugAge) | Barardo et al. 2017 | -0.23 | -0.36 | 92 |
| mice | IGF1 mutants | Garratt et al. 2017 | -0.37 | -0.48 | 42 |
| mice | mutant mice | de Magalhães et al. 2018 | NA | -0.55 | 33 |
| mice | UM-HET3 (NIA ITP) | Nadon et al. 2016; other | -0.27 | -0.33 | 395 |
| mice | various, on CR | Swindell et al. 2012 | 0.00 | -0.42 | 72 |
| rats | various, on CR | Swindell et al. 2012 | -0.33 | -0.56 | 53 |
|  |  | mean | -0.44 | -0.53 |  |

**Table S1. Correlation between control lifespan of mice and absolute change in lifespan, or relative change in lifespan, after various interventions.**
Data from Garratt et al. (2017) is for IGF1 mutants, see Fig. S11 for data on GH dwarfs. DrugAge data was curated and cleaned up for our purposes (e.g. excluding ITP and rapamycin data). We show correlation coefficients between control lifespan and lifespan extension of the treated group measured as fold-change (“rel” ) or measured as an absolute difference (“abs”). N refers to the number of groups or cohorts, not individual animals.

| **species** | **strain(s) / intervention** | **reference** | **N** | **LS ctrl** | **LS treated** |
| --- | --- | --- | --- | --- | --- |
| mice | rapamycin | Selvarani et al. 2021 | 30 | 811 | 930 |
| mice | telomerase induction | literature search | 8 | 773 | 914 |
| mice | comparative | Yuan et al. 2012 | 32 | 700 | NA |
| mice | comparative | Austad et al. 2011 | 118 | 710 | NA |
| mice | caloric restriction (CR) | literature search | 33 | 764 | 956 |
| mice | genetic interventions | Tacutu et al. 2018 | 24 | 843 | 1032 |

**Table S2. Other datasets used for comparative purposes**
We analyzed the **Austad et al. (2011)** dataset after excluding the **Yuan et al. (2012)** data which we analyze separately. Shown here is data for the full Austad dataset with male and female lifespans pooled. For the DrugAge dataset (**Tacutu et al. 2018**) we show only the top interventions we re-analyzed. N refers to the number of groups or cohorts, not individual animals. The lifespan (LS) of the control group and the treated group is shown in days.

| **species** | **strains (dataset)** | **reference** | **Pearson** | **Spearman** |
| --- | --- | --- | --- | --- |
| mice | metformin | Parish and Swindell 2022 | -0.49 | (-0.42) |
| yeast | deletion mutants | Schleit et al. 2013 | -0.24 | -0.28 |
| flies | DGRP inbred panel | Jin et al. 2020 | -0.38 | -0.34 |
| mice | ILSXISS inbred panel | Liao et al. 2010 | -0.58 | -0.51 |
| mice | ILSXISS inbred panel | Rikke et al. 2010 | -0.56 | -0.34 |
| mice | ILSXISS inbred panel | Unnikrishnan et al. 2021 | -0.71 | -0.64 |
| mice | Swiss mice | Schroeder et al. 1975 | -0.78 | -0.75 |
| mice | B/W and DBA/2f mice | Fernandes et al. 1976 | -0.68 | -0.53 |
| mice | various (DrugAge) | Barardo et al. 2017 | -0.23 | (-0.19) |
| mice | IGF1 mutants | Garratt et al. 2017 | -0.37 | -0.34 |
| mice | UM-HET3 (NIA ITP) | Nadon et al. 2016; other | -0.27 | -0.27 |
| mice | various, on CR | Swindell et al. 2012 | (0.00) | (0.03) |
| rats | various, on CR | Swindell et al. 2012 | -0.33 | (-0.25) |
|  |  | mean | -0.43 | -0.38 |

**Table S3**
Comparison of Pearson and Spearman correlation for the datasets in Table S1. Results that fail to reach statistical significance are shown in parentheses.

|  |  | **well controlled?** | |  |
| --- | --- | --- | --- | --- |
| **reference** | **study type** | **intervention** | **husbandry** | **R-value** |
| Liao et al. 2010 | CR experiment | +++ | +++ | -0.53 |
| Rikke et al. 2010 | CR experiment | +++ | +++ | -0.48 |
| Unnikrishnan et al. 2021 | CR experiment | +++ | +++ | -0.71 |
| Fernandes et al. 1976 | CR experiment | +++ | ++ | -0.68 |
| NIA ITP | feeding experiment | ++ | ++ | -0.27 |
| Schroeder et al. 1975 | feeding experiment | 0 | ++ | -0.78 |
| Swindell et al. 2012 | meta-analysis of CR | ++ | 0 | (0.00) |
| Parish and Swindell 2022 | meta-analysis | ++ | 0 | -0.49 |
| Garratt et al. 2017 | meta-analysis | + | 0 | -0.37 |
| DrugAge | meta-analysis | 0 | 0 | (-0.09) |
| de Magalhães et al. (2018) | meta-analysis | 0 | 0 | -0.55 |

**Table S4.**
Better controlled mouse studies or datasets show a more robust signal with a higher negative correlation between control lifespan and experimental lifespan extension. The term “well-controlled” here refers to the consistency of husbandry conditions or interventional treatment. Data from Garratt et al. (2017) is for IGF1 mutants. Correlations that fail to reach significance are in parenthesis. Intervention (+++): the same intervention tested across all conditions; intervention (++): the same or a similar intervention tested across all conditions (e.g. differences in the intensity of CR); intervention (+): similar intervention tested across all conditions (e.g. mutations of different genes in the same pathway). Husbandry (+++): animals tested under the same conditions in one experiment; husbandry (++): animals tested by the same lab in one or multiple studies published close together.

|  | female |  |  |  | male |  |  |
| --- | --- | --- | --- | --- | --- | --- | --- |
| site | **TJL** | **UM** | **UT** |  | **TJL** | **UM** | **UT** |
| R-value | -0.086 | -0.213^ | -0.431* |  | -0.400* | -0.226^ | -0.443* |
| p-value | 0.498 | 0.091 | 0.0004 |  | 0.0007 | 0.064 | 0.0002 |

**Table S5**
Correlation between control lifespan in the Interventions Testing Program and treatment effect (change in lifespan compared to control) across three testing sites and two genders. TJL=the Jackson Laboratory, UM=University of Michigan, UT=University of Texas Health Science Center. ^indicates p<0.10, *indicates p<0.05.

| t-test | TJL | UM | UT |  | log-rank | TJL | UM | UT |
| --- | --- | --- | --- | --- | --- | --- | --- | --- |
| male | 15 | 15 | 29 |  | male | 17 | 14 | 27 |
| t-test | **TJL** | **UM** | **UT** |  | log-rank | **TJL** | **UM** | **UT** |
| female | 13 | 11 | 18 |  | female | 15 | 13 | 22 |

**Table S6. How common were significant lifespan effects across the three sites?**
This table shows how many treatments significantly affected mouse lifespan by t-test and log-rank test across the three testing sites of the interventions testing program. Data is stratified by testing site and gender. TJL=the Jackson Laboratory, UM=University of Michigan, UT=University of Texas Health Science Center.

|  | **male** | **female** | **female** |
| --- | --- | --- | --- |
|  | ILSXISS | ILSXISS | excl. Rikke et al. |
| **median** | 938 | 835 | 897 |
| **mean** | 919 | 846 | 881 |
| **top Q** | 1051 | 961 | 1018 |

**Table S7. Summary statistics for different ILSXISS mouse strains**
We present here median, mean and 20^th^ percentile lifespans pooled from three studies using ILSXISS mice (**Liao et al. 2010, Rikke et al. 2010,** **Unnikrishnan et al. 2021**). Since the study by Rikke et al. 2010 reported lower lifespans for the same strains tested by Liao et al. 2010 we also show data after excluding this study.

|  | sex | mean | median | top_q |
| --- | --- | --- | --- | --- |
| B6 | F | 721 | 736 | 818 |
| B6 | M | 738 | 778 | 842 |
| ILSXISS | F | 835 | 846 | 961 |
| ILSXISS | M | 919 | 938 | 1051 |
| Yuan | F | 713 | 724 | 808 |
| Yuan | M | 713 | 718 | 837 |
| UM-HET3 | F | 882 | 883 | 899 |
| UM-HET3 | M | 798 | 800 | 862 |
| mean |  | 790 | 803 | 885 |

**Table S8. Summary statistics for different mouse strains**
We present here median, mean and 20^th^ percentile lifespans for the strains plotted in Fig. 4.

|  | ITP (sig. LS extension?) | | DrugAge (900-day rule) | |
| --- | --- | --- | --- | --- |
| **Intervention (ITP)** | **males** | **females** | **LS - pass** | **LS - fail** |
| NDGA | 4/4 | 0/4 | 2 | 2 |
| rapamycin | 7/8 | 7/8 | 6 | 7 |
| aspirin | 1/3 | 0/3 | NA | 3 |
| NR* | 0/1 | 0/1 | NA | 3 |
| metformin | 0/1 | 0/1 | NA | 4 |
| curcumin | 0/1 | 0/1 | NA | 2 |
| green tea extract | 0/1 | 0/1 | NA | 1 |

**Table S9. Translation between DrugAge and interventions testing program (ITP)**
When counting whether an intervention passes the 900-day rule we consider each individual dataset as reported in DrugAge. Success in the ITP is defined as significant lifespan (LS) extension in the pooled male or female dataset. Compounds were matched using reasonable assumptions, e.g. nicotinamide riboside (NR) was tested in the ITP and we compare it to NR and nicotinamide as reported in DrugAge.

| **ID** | **shorthand** | **drug name** | **count** |
| --- | --- | --- | --- |
| 1 | ACA_16 | Acarbose | 1 |
| 2 | ACA_mid | Acarbose | 2 |
| 3 | BD | 1,3-butanediol | 1 |
| 4 | Cape_hi | Caffeic acid phenethyl ester | 1 |
| 5 | Capt | Captopril | 1 |
| 6 | Est_hi | 17-α-estradiol | 1 |
| 7 | Gly | Glycine | 1 |
| 8 | MB | Methylene Blue | 1 |
| 9 | MetRapa | Rapamycin, Metformin | 4 |
| 10 | NR | Nicotinamide Riboside | 1 |
| 11 | R09_hi | Rapamycin | 4 |
| 12 | R09_lo | Rapamycin | 4 |
| 13 | R09_mid | Rapamycin | 3 |
| 14 | RaAc16 | Rapamycin, Acarbose | 1 |
| 15 | RaAc9 | Rapamycin, Acarbose | 2 |
| 16 | Rapa_20hi | Rapamycin | 2 |
| 17 | Rapa_onoff | Rapamycin | 2 |
| 18 | Rapa_Y2 | Rapamycin | 4 |
| 19 | Rapa_Y3 | Rapamycin | 3 |

**Table S10. Drugs in the Interventions Testing Program that meet the 900-day rule criteria in at least one cohort**
The table shows compounds meeting the 900-day rule criteria in any cohort of the ITP. We use the following definition of the rule: control lifespan ≥850 days and significant lifespan extension ≥50 days. Count indicates how many cohorts met the criteria with a maximum of 6 per compound in most cases since two genders were tested across three study sites. Compounds that extended lifespan in more than one cohort: acarbose, rapamycin, rapamycin-acarbose, rapamycin-metformin.

| **ID** | **shorthand** | **drug name** | **count** | **gender** |
| --- | --- | --- | --- | --- |
| 1 | MetRapa | Rapamycin, Metformin | 1 | F |
| 2 | R09_hi | Rapamycin | 1 | F |
| 3 | R09_lo | Rapamycin | 1 | F |
| 4 | R09_mid | Rapamycin | 1 | F |
| 5 | RaAc16 | Rapamycin, Acarbose | 1 | F |
| 6 | RaAc9 | Rapamycin, Acarbose | 1 | F |
| 7 | Rapa_20hi | Rapamycin | 1 | F |
| 8 | Rapa_onoff | Rapamycin | 1 | F |
| 9 | Rapa_Y2 | Rapamycin | 1 | F |
| 10 | Rapa_Y3 | Rapamycin | 1 | F |

**Table S11.** **Drugs in the Interventions Testing Program that meet the the 900-day rule criteria in the pooled dataset**
Compounds meeting the 900-day rule criteria in the ITP. We use the following definition of the rule: control LS ≥850 days and significant LS extension ≥50 days. Count indicates how many cohorts met the criteria with a maximum of 2 per compound in most cases since two genders were tested.

| **ID** | **names** | **name** | **count** |
| --- | --- | --- | --- |
| 1 | ACA | Acarbose | 3 |
| 2 | ACA_16 | Acarbose | 1 |
| 3 | ACA_hi | Acarbose | 3 |
| 4 | ACA_lo | Acarbose | 1 |
| 5 | ACA_mid | Acarbose | 3 |
| 6 | Cana | Canagliflozin | 2 |
| 7 | Capt | Captopril | 1 |
| 8 | Enal | Enalapril | 1 |
| 9 | Est | 17-α-estradiol | 1 |
| 10 | Est_16 | 17-α-estradiol | 3 |
| 11 | Est_20 | 17-α-estradiol | 1 |
| 12 | Est_hi | 17-α-estradiol | 3 |
| 13 | MB | Methylene blue | 1 |
| 14 | MetRapa | Rapamycin, Metformin | 5 |
| 15 | ND_lo | Nordihydroguaiaretic acid | 1 |
| 16 | NDGA | Nordihydroguaiaretic acid | 1 |
| 17 | NR | Nicotinamide Riboside | 1 |
| 18 | R09_hi | Rapamycin | 5 |
| 19 | R09_lo | Rapamycin | 3 |
| 20 | R09_mid | Rapamycin | 5 |
| 21 | RaAc16 | Rapamycin, Acarbose | 3 |
| 22 | RaAc9 | Rapamycin, Acarbose | 5 |
| 23 | Rapa_20hi | Rapamycin | 3 |
| 24 | Rapa_onoff | Rapamycin | 2 |
| 25 | Rapa_stop | Rapamycin | 2 |
| 26 | Rapa_Y2 | Rapamycin | 4 |
| 27 | Rapa_Y3 | Rapamycin | 4 |
| 28 | Res_hi3 | Resveratrol | 1 |
| 29 | UA | Ursolic acid | 1 |

**Table S12. Drugs in the Interventions Testing Program showing larger than expected lifespan extension**
Compounds that led to lifespan (LS)extension of >50 days above the predicted LS extension in in at least one cohort. Count indicates how many cohorts met the criteria (maximum of 6 per compound tested in both genders). Compounds that extended lifespan in more than one cohort: Acarbose, Canagliflozin, 17-α-estradiol, Nordihydroguaiaretic acid, Rapamycin, Rapamycin combinations and ACE inhibitors.

|  |  |  |  |  | **dose (ppm)** | |  |
| --- | --- | --- | --- | --- | --- | --- | --- |
| **ID** | **names** | **name** | **LS residual (d)** | **year** | **Rapa** | **Compound 2** | **Start (mo)** |
| 1 | RaAc9 | Rapa+Acarbose | 158.3 | 2017 | 14 | 1000 | 9 |
| 2 | R09_hi | Rapamycin | 147.8 | 2009 | 42 | NA | 9 |
| 3 | MetRapa | Rapa+Metformin | 147.4 | 2011 | 14 | 1000 | 9 |
| 4 | Est_16 | 17-α-estradiol | 98.5 | 2016 | NA | 14.4 | 16 |
| 5 | Rapa_Y2 | Rapamycin | 92.2 | 2005 | 14 | NA | 20 |
| 6 | R09_mid | Rapamycin | 76.0 | 2009 | 14 | NA | 9 |
| 7 | Rapa_Y3 | Rapamycin | 75.3 | 2006 | 14 | NA | 9 |
| 8 | ACA | Acarbose | 74.9 | 2009 | NA | 1000 | 4 |
| 9 | Rapa_20hi | Rapamycin | 60.0 | 2015 | 42 | NA | 20-23 |
| 10 | ACA_mid | Acarbose | 57.2 | 2013.0 | NA | 1000 | 8 |

**Table S13. Top 10 interventions with larger than expected lifespan extension**
The top 10 interventions ranked by the lifespan (LS) extension^actual-predicted^ (i.e. lifespan residual in days). In the “Rapa_20hi” group rapamycin (Rapa) was only given between 20-23 months (mo) of age.

|  | **compound_name** | **strain** | **gender** | **pubmed_id** | **LS.intervn** | **LS.ctrl** |
| --- | --- | --- | --- | --- | --- | --- |
| 1 | Nordihydroguaiaretic acid | B6C3F1 | male | 25380600 | 1090 | 980 |
| 2 | Metoprolol | C3B6F1 | male | 23314750 | 1080 | 983 |
| 3 | L-deprenyl | B6CBAF1 | male | 8914495 | 1073 | 1003 |
| 4 | Nordihydroguaiaretic acid | B6C3F1 | male | 25380600 | 1052 | 980 |
| 5 | Nebivolol | C3B6F1 | male | 23314750 | 1046 | 983 |
| 6 | β-Aminopropionitrile fumarate | LAF/J | male | 689120 | 1041 | 984 |
| 7 | β-Aminopropionitrile fumarate | LAF/J | male | 689120 | 1038 | 984 |
| 8 | Polyphenols | C57BL/6J | male | 22265987 | 1005 | 945 |
| 9 | N-acetyl-L-cysteine | UM-HET3 | male | 20819793 | 1000 | 831 |
| 10 | Spermidine | C57BL/6 | male | 28386016 | 999 | 807 |
| 11 | Dasatinib + Quercetin | C57BL/6 | combined | 29988130 | 996 | 937 |
| 12 | L2-Cmu (IGF-1R mAb) | CB6F1 | female | 29921922 | 966 | 847 |
| 13 | Ivabradine | C57BL/6J | male | 25589054 | 962 | 906 |
| 14 | CASIN (Cdc42 inhibitor) | C57BL/6 | female | 32755011 | 950 | 868 |

**Table S14. Top lifespan extending compounds in DrugAge**
In total, 14 datasets featuring 12 unique compounds met the 900-day rule among compounds reported in DrugAge (Barardo et al. 2017). We only included compounds if they produced LS extension >50 days, compared to control animals with a LS of ≥850 days, or resulted in a treated LS of ≥950 days.

| **Nr** | **Reference** | **Gene** | **type** | **putative pathway(s)** |
| --- | --- | --- | --- | --- |
| 1 | Brown-Borg et al. 1996 | Prop1 | KO | GH/IGF-1/Insulin pathway |
| 2 | Zhang et al. 2012 | Fgf-21 | Tg | growth/nutrient signalling |
| 3 | Flurkey et al. 2001 | Ghrhr | KO | GH/IGF-1/Insulin pathway |
| 4 | Flurkey et al. 2001 | Pou1f1 | KO | GH/IGF-1/Insulin pathway |
| 5 | Ren et al. 2017 | Akt-2 | KO | growth/nutrient signalling |
| 6 | Tomas-Loba et al. 2008 | Tert | Tg | other |
| 7 | Kanfi et al. 2012 | Sirt6 | Tg | other |
| 8 | Riera et al. 2014 | Trpv1 | KO | other |
| 9 | Ran et al. 2007 | Gpx4 | KO (+/-) | other |
| 10 | Coschigano et al. 2003 | Ghr | KO | GH/IGF-1/Insulin pathway |
| 11 | Liu et al. 2005 | Clk-1 | KO (+/-) | other |
| 12 | Enns et al. 2009 | Prkar2b | KO | other |
| 13 | Blueher et al. 2003 | Insr | KO, adipose | GH/IGF-1/Insulin pathway |
| 14 | Canaan et al. 2014 | Ubd | KO | other |
| 15 | Kurosu et al. 2005 | Kl | Tg | GH/IGF-1/Insulin pathway |
| 16 | Yan et al. 2007 | Ac5 | KO | other |
| 17 | López-Andrés 2013 | Ct-1 | KO | other |
| 18 | Hofmann et al. 2015 | Myc | KO (+/-) | growth/nutrient signalling |
| 19 | Selman et al. 2016 | Rps6kb1 | KO | mTOR, translation |
| 20 | Selman et al. 2008 | Irs1 | KO | GH/IGF-1/Insulin pathway |
| 21 | Wu et al. 2012 | Cisd2 | Tg | other |
| 22 | Zhang et al. 2013 | Ikbkb | viral, KO | other |
| 23 | Tucker et al. 2008 | Kcna3 | KO | other |
| 24 | Wu et al. 2013 | mTor | hypomorphic | mTOR pathway |

**Table S15.**
We identified 24 genes from GenAge that meet the criteria of the 900-day rule, i.e. extend lifespan in long-lived controls or lead to LS ≥950 days. Since the precise mechanisms of action for most of these genes are not known we group them roughly into four categories: mTOR signalling, growth signalling, GH/IGF-1/Insulin-axis and other pathways.

| **Intervention** | **meets 900-day rule?** | **LS extension in long lived mice?** |
| --- | --- | --- |
| Telomerase | 3 out of 9 (38%) | 2 out of 2 (100%) |
| Rapamycin | 16 out of 30 (53%) | 14 out of 15 (93%) |
| Metformin | 1 out of 20 (5%) | 2 out of 8 (25%) |

**Table S16.**
We use the following definition of the 900-day rule: control lifespan (LS) ≥850 days and LS extension ≥50 days or treated LS ≥950 days. Long-lived mice are defined as those with an LS of ≥850 days.

**Supplementary results and discussion**

**Pituitary dwarfism highlights the usefulness of the 900-day rule**
Early studies reported pituitary dwarf mice to be short-lived and their lifespan was extended by growth hormone and thyroxine treatment. In this study controls lived to 600 days and dwarfs to 150 days (**Fabris et al. 1972**). In contrast, **Brown-Borg (1996)** were the first to report longer lifespans of Ames pituitary dwarfs. Although control mice in this study were also short-lived with lifespans around 700 days, the dwarf mice lived to over 1000 days. According to the 900 day-rule we should have higher confidence in the latter result than the former.

Since then, gradually a consensus has emerged that Ames, Snell and GRKO dwarfism all robustly extend mouse lifespan. In a reanalysis of the meta-analysis by **Garratt et al. (2017)**, 4 out of 17 control groups in dwarf studies were long-lived passing the 900-day rule. In all of these, dwarfism extended lifespan. Perhaps more importantly, even though controls were short-lived in several other studies reported, the treated groups were exceptionally long-lived, which highlights the importance of comparing treated animals with historical controls in such cases (**Fig. S21**). In 11 out of 17 studies dwarf mice lived over 950 days beating the benchmark for historical inbred controls (900 ± 50 days).
